# Genetic determinants of intraspecific variation in crossover frequencies in the honeybee, *Apis mellifera*

**DOI:** 10.64898/2026.02.03.702590

**Authors:** Turid Everitt, Luong Hieu Trinh, Demetris Taliadoros, Tilman Rönneburg, Anna Olsson, Joachim R. de Miranda, Bertrand Servin, Matthew T. Webster

## Abstract

Meiotic recombination facilitates natural selection and is necessary for correct chromosomal segregation in most sexually reproducing species. Crossover rates vary greatly both within and among species, but the determinants of this variation are not fully understood. The honeybee *Apis mellifera* has extremely high recombination rates. Honeybee males (drones) are haploid, which enables the distribution of crossovers to be directly estimated from the progeny of a single reproductive female (queen). Here we map crossover events in the honeybee using whole genome sequencing of 1509 drone progeny of 184 queens. This allows us to assay intra-specific variation in recombination rate and its genetic and non-genetic determinants. We estimate the average crossover rate as 23 cM/Mb, with between 22 and 88 crossovers events detected in individual offspring. We estimate 28% of this variation is additive heritable variation among queens. There is no effect of queen age or genetic background on crossover rate. A genome-wide association study identifies variation in the gene *mlh1* as associated with mean crossover rate. We estimate that variation in the gene is associated with a 10% difference in crossover rate between the two homozygous genotypes at the most significant SNP. This gene has a well-established role in recombination and variation in the gene could affect crossover rates by affecting resolution of Holliday junctions as crossovers. This is the first gene discovered to be associated with recombination rate variation in an insect. Adaptive evolution of this gene could potentially underlie the extremely high recombination rates in honeybees.

## Introduction

Homologous recombination is an essential biological process that occurs during meiosis in sexually reproducing organisms. It is initiated at chromosomal double strand breaks (DSBs) and leads to either a reciprocal exchange of genetic material between homologous chromosomes (crossover, CO) or copying of a shorter segment of one chromosome to the other (non-crossover, NCO). It is commonly accepted that the advantage of recombination in sexual taxa is that it breaks up associations between advantageous and disadvantageous alleles at linked loci allowing selection to act on them independently (Hill and Robertson 1966; Hartfield and Keightley 2012). This facilitates beneficial variants to become fixed and deleterious ones to become purged from populations. The increased haplotype diversity created through recombination could be particularly beneficial for adaptation to changing environmental conditions and resistance to parasites (Hamilton et al. 1990; Hartfield and Keightley 2012).

Recombination is also important for maintaining genetic integrity during meiosis through the formation of chiasmata, which are required for correct segregation of homologous chromosomes. In most species, proper chromosomal segregation seems to require at least one CO event per bivalent (Jones 1984; Berchowitz and Copenhaver 2010). Abnormal recombination increases the risk of nondisjunction and aneuploidies (Koehler et al. 1996; Hassold and Hunt 2001) and can lead to chromosomal rearrangements via non-allelic homologous recombination. Some studies also indicate increased risk of *de novo* mutations near DSBs in humans (Halldorsson et al. 2019; Hinch et al. 2023).

Despite its essential functions, the frequency at which recombination occurs along chromosomes varies over multiple orders of magnitude among species of sexually reproducing eukaryotes (Wilfert et al. 2007; Stapley et al. 2017). These rates are likely determined by a balance between the costs and benefits of meiotic recombination, which are influenced by both molecular and evolutionary constraints. Our understanding of how these different factors constrain the recombination rate in different groups of organisms is however far from complete.

Recombination rate also varies between individuals of the same species. An important way to understand the forces governing evolution of recombination rate is to examine the causes of this inter-individual variation and in particular its underlying genetic architecture. Studies of inter-individual variation in recombination rate have mainly focussed on humans and domestic vertebrates, although studies of wild animals, plants, fungi and invertebrates have also been performed. These studies have revealed the presence of substantially heritable variation in CO rates in many taxa (Johnston 2024). For example, the narrow-sense heritability of recombination rate has been estimated as 0.46 in mice (Dumont et al. 2009), 0.22 in cattle (Sandor et al. 2012) and 0.12 - 0.16 in Soay sheep (Johnston et al. 2016).

Specific major effect loci underlying variation in CO frequencies can be identified through genome-wide association studies (GWAS). Several studies in a large number of mammalian taxa have identified major effect loci with strong effects on the genome-wide recombination rate. The majority of genes identified have clear roles in meiotic process, including the formation of double-stranded breaks, synapsis, CO/NCO decision and CO resolution (Dapper and Payseur 2019; Johnston 2024). In house sparrows and Atlantic salmon, recombination rate seems to have a more polygenic basis with a great number of low effect loci involved (Brekke, Johnston, Knutsen, et al. 2023; Johnston 2024; McAuley et al. 2024). In *Drosophila melanogaster*, a number of genes were found to affect the recombination rate, none of which are included in known pathways for meiotic recombination (Hunter, Huang, et al. 2016). The genetic control of recombination rate in other insects is not well studied and many aspects remain to be investigated.

Recombination rates also exhibit plasticity, and have been shown to be influenced by multiple extrinsic and intrinsic factors (Modliszewski and Copenhaver 2017). Extrinsic factors such as temperature, nutrient availability or desiccation have been found to alter the recombination rate in species of insects, mammals and plants (Rybnikov et al. 2023). *Drosophila* parasite infection increases recombination rate (Singh et al. 2015). While it seems clear that stress from the environment can alter recombination rates, whether a particular stressor increases or decreases recombination is not consistent between species or stressors. Age is another factor shown to correlate with recombination rate. In humans, there is a weak but significant association between high maternal age and elevated recombination rate in the mother (Campbell et al. 2015; Martin et al. 2015). The cause of this relationship is unclear and must be reconciled with the fact that recombination occurs during fetal development in human females. A maternal age effect has been found to affect the recombination rate in other taxa of animals and plants, but the type of effect is not consistent as both increasing and decreasing rates with age have been reported (Polani and Jagiello 1976; Hussin et al. 2011; Tedman-Aucoin and Agrawal 2012; Hunter, Robinson, et al. 2016; Lozada-Soto et al. 2021; Shen et al. 2021).

Social insects from the order Hymenoptera have considerably higher recombination rates than other non-social insects and most other metazoan taxa (Wilfert et al. 2007). The highest rates are found in honey bees from the genus *Apis* that have recombination rates in the range of 19 - 37 cM/Mb (Shi et al. 2013; Wallberg et al. 2015; Rueppell et al. 2016; Jones et al. 2019; Kawakami et al. 2019). This suggests natural selection has acted to increase the recombination rate in these taxa. Elevated recombination rates appear to be a feature of social insects in the Hymenoptera order specifically, as termites, which are social insects from the order Blattodea, do not have elevated recombination rates (Everitt et al. 2025). There have been multiple hypotheses trying to explain this finding, and understanding the evolutionary forces driving high recombination rates in social Hymenoptera could help us to understand the forces that determine recombination rates in general.

All species of Hymenoptera are haplodiploid. Females are diploid, whereas males are haploid and develop from unfertilized eggs. Recombination only occurs in reproductively-capable females, and direct sequencing of their haploid (male) offspring can be used to infer the location of recombination events. This method has been applied previously to honeybees by (Liu et al. 2015; Kawakami et al. 2019) confirming recombination rates in the vicinity of 25 cM/Mb. In addition it has been used to analyse inter-individual variation in recombination rate between females. Kawakami *et al*. analysed genome sequences from 299 haploid offspring from 22 colonies of honeybees that varied in genetic background and 9 colonies of bumblebees. This study identified substantial variation between individuals, to the extent that the distribution of CO frequencies among colonies overlapped between species, despite a greater than twofold difference in the mean. This study also estimated broad sense heritability to be 44% in the honey bee (Kawakami et al. 2019). In addition, a significant genetic background effect was observed, with honeybees from African subspecies exhibiting lower average CO rates than European subspecies.

Honeybees are an ideal study system for analysing variation in recombination rate and the genetic and non-genetic factors underlying it. So far there have been few studies investigating the genetic basis of recombination rate variation in insects, and no major-effect loci have been identified. Honeybees are particularly amenable for studying recombination as CO events can be directly inferred using the genome sequence of haploid males that are identical to female gametes. Each honeybee colony is headed by a single reproductive female queen, which can live for 3-4 years and continue to produce a great number of offspring throughout its life. This is an especially suitable system for studying the effect of maternal age on recombination rate.

Here we generate fine-scale maps of CO events across the honey bee genome using whole-genome sequencing of 1509 drone progeny of 184 queens. The results allow us to analyse the causes of individual variation in CO frequencies. Our main aims were 1) to analyse the effect of maternal age and genetic background on CO frequencies, 2) to estimate the proportion of variation in CO rate that is heritable and 3) to identify specific genes with a major effect on variation in CO rate.

## Results

### Substantial variation in recombination rate between individuals

We analysed a dataset of whole-genome sequencing data of 1509 honey bee drones from 184 different colonies, consisting of previously published sequencing data of drones from 16 colonies (Kawakami et al. 2019) and newly-sequenced samples (Supplementary Tables S1,S2). All sequences were mapped to the Amel_HAv3.1 reference genome (Wallberg et al. 2019), followed by variant calling and filtering. For the purpose of inferring the position of COs in each drone, we generated a strictly-filtered dataset of common high-confidence SNPs. This filtered dataset has an average read depth of 7.5 x per sample and contains 326,130 SNPs from 1509 honey bee drones and 184 different colonies, with eight drones per colony in most cases (Supplementary Table S1). The relatedness between individuals was measured by the *A_jk_* score as defined by (Yang et al. 2010) and the closest relatives of all samples were confirmed to be from the same colony (Supplementary Figure S1 A-D; Supplementary Table S3).

Using the software YAPP (Servin 2025), we first reconstructed the phased, diploid sequence of the queen in each colony and then inferred recombination events in each drone where the drone haplotype switched between the two queen haplotypes, as illustrated by an example colony in Figure 1. In this way, we could accurately map the position of COs and obtain detailed recombination rate estimates for each meiosis. The CO positions are identified in terms of intervals within which the haplotype switches and the mean length of those intervals was 250 kbp, with >75% of the intervals shorter than 300 kbp. In total we identified 74,905 COs in the set of 1509 drones (Figure 2 A,B). The mean number of COs per drone is 50, with a range between 22 and 88.

**Figure 1.**
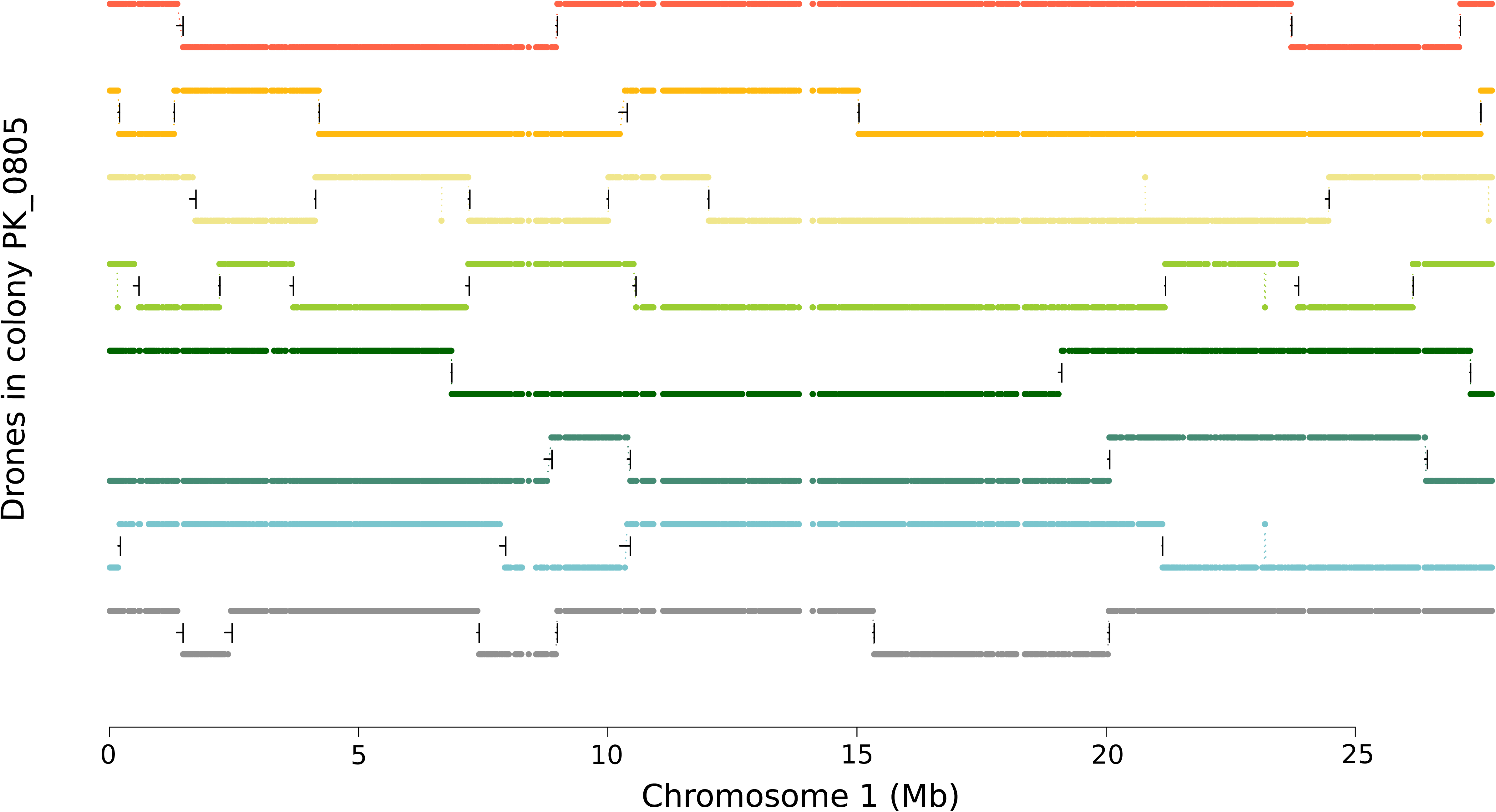
Haplotypes of drones from one colony on chromosome 1, with each color representing one drone. Each drone can have two different values on the y-axis, representing the two queen haplotypes and showing from which of those the drone most likely originates at each chromosomal position. When a drone switches between its two values on the y-axis, this indicates a recombination event. The intervals marked with black lines are the intervals within which a recombination event was identified by the software YAPP (a horizontal line between the start and end of the interval and a vertical line at the end of the interval).

**Figure 2.**
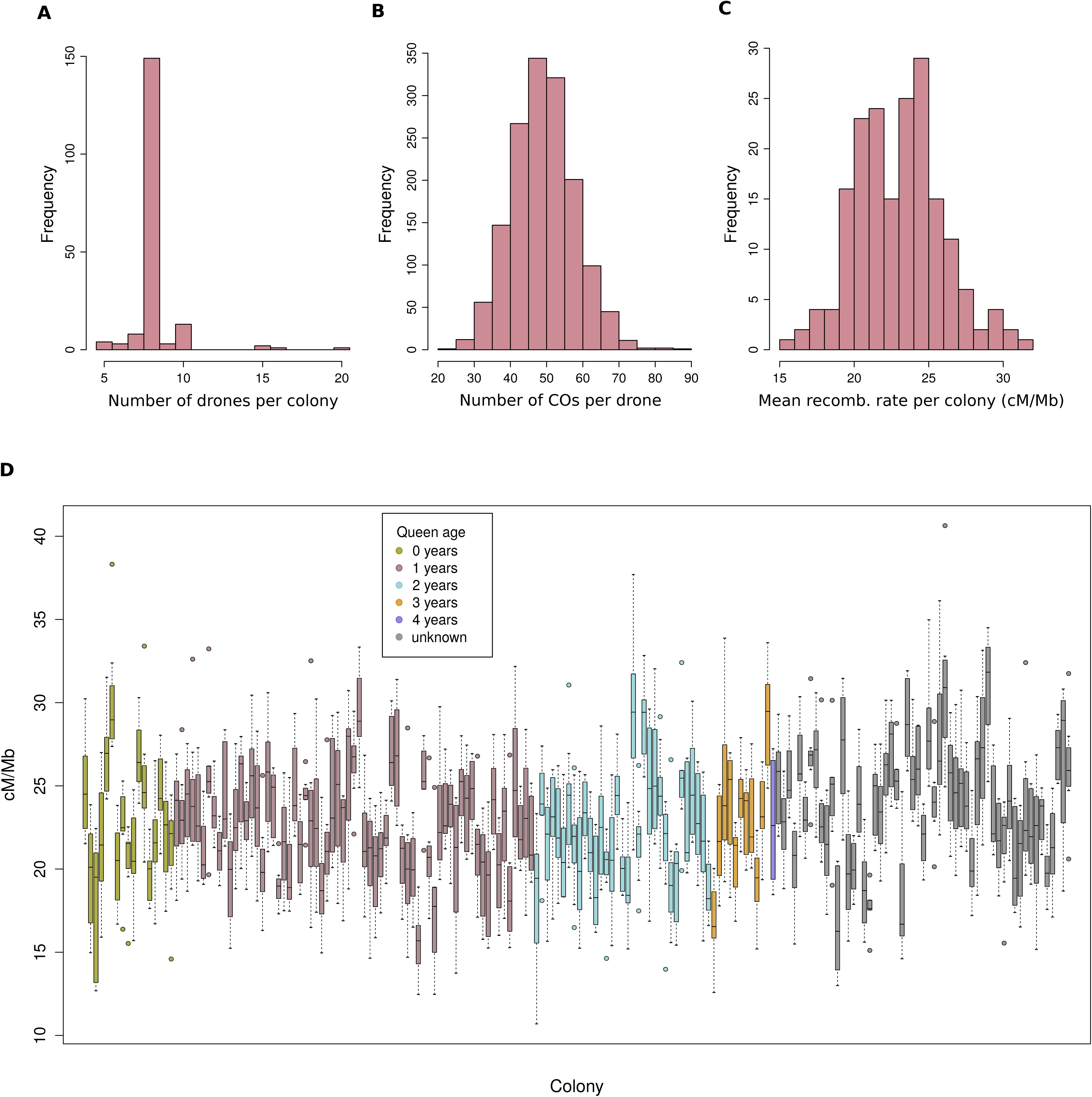
Variation in crossover frequencies in the dataset of 1509 drones derived from 184 queens. **A)** distribution of number of drones per colony in the dataset, **B)** distribution of total number of crossovers inferred per drone, **C)** distribution of mean crossover rate per colony. **D)** Boxplot of crossover frequency per drone, divided by colony (queen). Each box represents one colony and is colored by the age of the queen.

Seventeen of the drone samples were sequenced twice and the replicates were treated as separate individuals for mapping, variant calling and recombination inference. Even though they were processed independently, all replicate samples yielded similar results from the recombination inference with YAPP (out of the 808 unique COs identified in total, 712 COs had the exact same coordinates in both replicates, 92 COs had partly overlapping intervals and only four COs did not match any other CO in the corresponding replicate; Supplementary Figure S2). This proves the robustness and reliability of the inference method.

The average genome wide recombination rate of all meioses was estimated as 23.04 cM/Mb with a standard deviation of 3.96 cM/Mb, which is similar to previous estimates of 26 cM/Mb (Wallberg et al. 2015; Kawakami et al. 2019) and 24 cM/Mb (Jones et al. 2019). There is substantial variation in the population-averaged recombination rate along chromosomes (Supplementary Figure S3; Supplementary Table S4) with regions of low recombination within previously identified pericentromeric regions (Wallberg et al. 2019; Everitt et al. 2023).

As can be seen in Figure 2 C,D, there is also considerable variation in recombination rate between colonies as well as between individual meioses (drones). This variation might be caused by both genetic and non-genetic factors, which we evaluate below in terms of the heritability, the effect of the genomic background, the effect of the queen age and a genome-wide association study to identify major-effect loci.

### High heritability of CO frequency and low heritability of allelic shuffling

The overall effect of genetic contributions to the recombination rate variability was estimated as the heritability in two different ways. A comparison of the recombination rate variance between colonies to the variance within each colony, based on a one-way ANOVA (Table 1), gave a broad-sense heritability *H^2^* = 52%. We also estimated both the broad- and narrow sense heritabilities based on a restricted maximum likelihood (ReML) model, with a genomic relationship matrix (GRM) as a covariant of the genetic effects. With this method, the broad-sense heritability *H*^2^ was estimated to 47% and the narrow-sense heritability *h*^2^ to 28%. The broad-sense heritability estimates are similar to the previous estimate of *H*^2^ = 44% in honeybees (Kawakami et al. 2019), while the narrow-sense heritability, which specifically captures only the additive genetic effects, is lower but not close to zero. These results suggest that a substantial proportion of the recombination rate variability is explained by genetics.

**Table 1.** One-way ANOVA of recombination rate per drone with colony as a random effect factor.

|  | DF | Sum Sq | Mean Sq | F-value | *p*-value |
| --- | --- | --- | --- | --- | --- |
| Colony | 183 | 1.24E-12 | 6.79E-15 | 7.9831 | < 2.2e-16 |
| Residuals | 1325 | 1.13E-12 | 8.51E-16 |  |  |

We used the number and locations of COs in the genome of each drone to estimate the intra-chromosomal allelic shuffling rate per meiosis, i.e. the probability that two randomly selected loci on the same chromosome have different parental origins, as defined by (Veller et al. 2019). The Spearman correlation between this measure and the CO count per drone is highly significant with *ρ* = 0.43 and *p* < 2.2*10^-16^ (Figure 3). We used ReML to estimate the heritability of shuffling, giving *H^2^* = 12% and *h^2^* = 6%, both values lower than the corresponding heritability estimates for CO frequency. When correcting for CO frequency, the heritability of shuffling is further reduced (*H^2^* = 5% and *h^2^*= 0.3%). Hence, out of these two correlated traits, CO frequency seems to be under stronger genetic control than intra-chromosomal shuffling.

**Figure 3.**
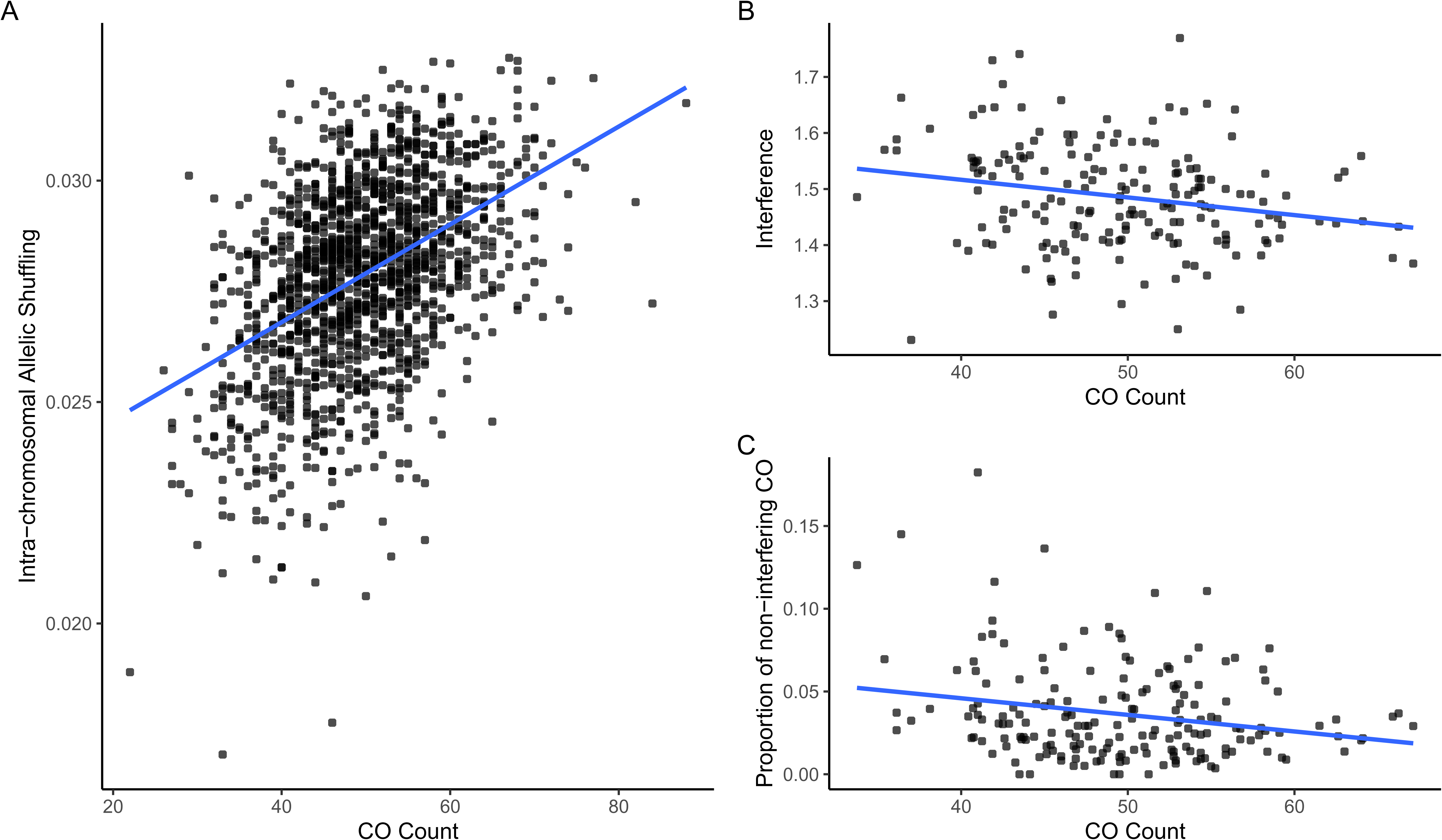
Correlation between CO count and recombination traits in queens and drones. A) CO count versus intra-chromosomal allelic shuffling per drone, Spearman’s *⍴* = 0.43, *p*-value 2.2e-16. B) CO count versus the strength of crossover interference per queen, Spearman’s *⍴* = -0.23, *p*-value = 0.0013. C) CO count versus the proportion of non-interfering COs per queen, Spearman’s *⍴* = -0.16, *p*-value = 0.033.

### Interference does not have a big effect on the CO distribution

We estimated the strength of CO interference at the colony level using the gamma model (Broman and Weber 2000). We found that CO interference is low in honeybees (mean 𝜈 = 1.5), which is consistent with previous results from (Kawakami et al. 2019) and can be compared to 𝜈 > 3 in bumblebees and most mammals and plants (Kawakami et al. 2019; Otto and Payseur 2019). The proportion of non-interfering COs is <5% for most colonies (mean 3.5%; Figure 3). The proportion estimates are similar to values in mammals and plants and therefore do not seem to be related to the low interference strength of honeybees (Basu-Roy et al. 2013; Campbell et al. 2016; Wang et al. 2016; Otto and Payseur 2019). Both the strength of the CO interference and the proportion of non-interfering COs have weak but significant negative correlations to the CO count (Spearman’s *ρ* = -0.23, *p* = 0.0013 for interference strength; Spearman’s *ρ* = -0.16, *p* = 0.033 for proportion of non-interfering COs; Figure 3). The two interference related traits are however not correlated to each other (Spearman’s *ρ* = -0.046, *p* = 0.53). The results indicate that although most COs in honey bees are from the interfering class (class I), which is the most common type in animals and plants, the degree of CO interference is low, which could be associated with the high number of COs on each honey bee chromosome (1.6 - 5.9 on average per chromosome).

### Crossover rate is not associated to queen age or genetic background

In order to evaluate the effect of the genetic background on the recombination rate, genetic variation in the drone population was analysed with principal component analysis (PCA). Two additional reference populations were included for comparison, based on the definitions by (Wallberg et al. 2014): the C-lineage (Harpur et al. 2014) and the M-lineage (Henriques et al. 2018). In the PCA (Figure 4A), the drones in each colony cluster together as expected, and the C and M reference populations are at the upper and lower ends, respectively, of the first component. The colonies appear between the reference populations, indicating that they are a mixture of the C- and M-lineages, which is typical for honey bees in this part of Europe (Wallberg et al. 2014). Most colonies are closer to the C-lineage reference population and those also have a greater spread along the second principal component.

**Figure 4.**
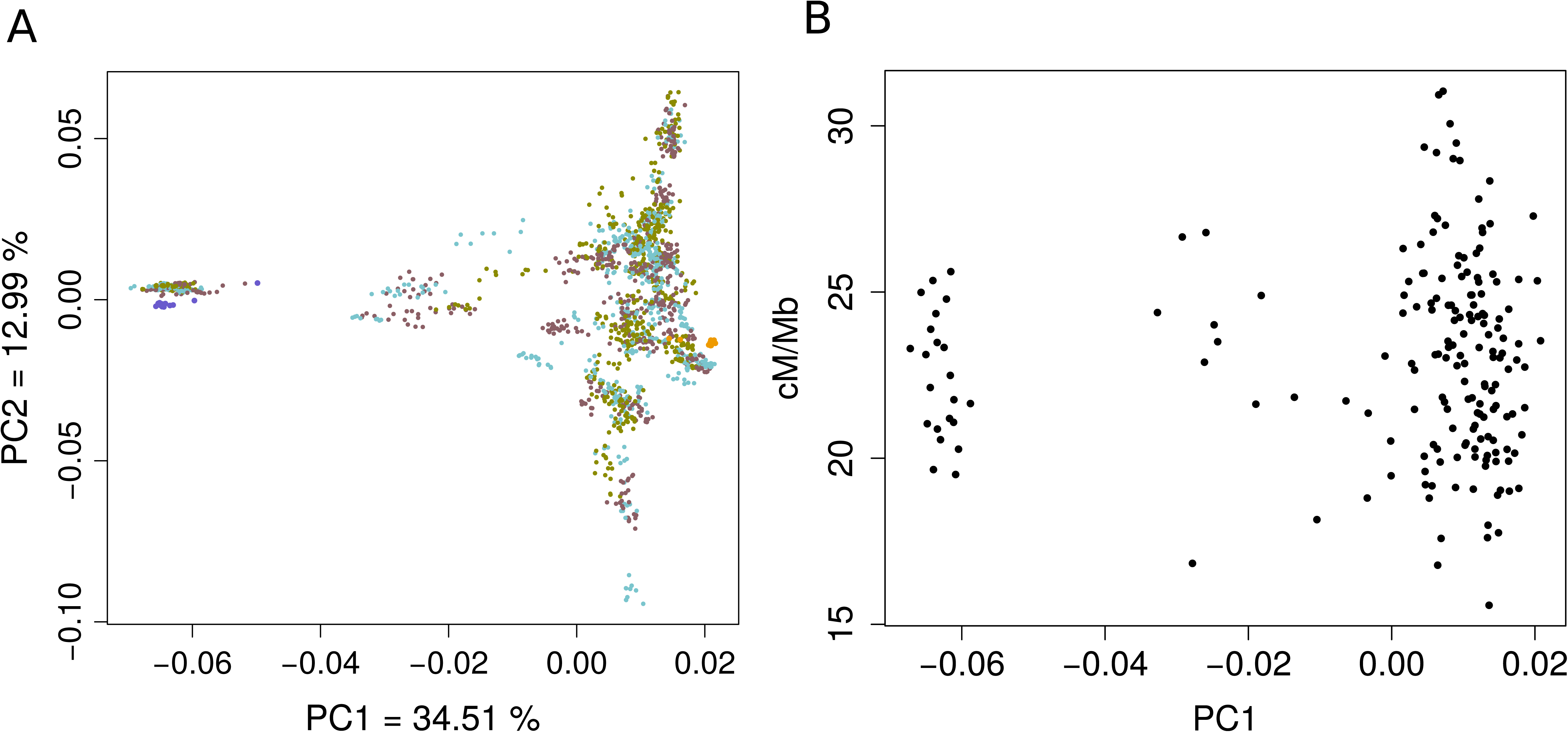
Effect of population structure on mean crossover rate. **A)** PCA of drones colored by colony (brown, light blue and green, repeated) and the reference populations from the C-lineage (yellow) and the M-lineage (blue). **B)** Crossover rate versus the first principal component from the PCA, mean values per colony.

The effect of the genomic background on the recombination rate was evaluated as the Spearman correlation between the first principal component of the PCA and the mean recombination rate per colony (Figure 4B). This gave a two-tailed *p*-value of 0.3 and 𝜌 = -0.076, showing that the genomic background does not have a significant effect on the recombination rate.

The effect of the maternal age was evaluated based on the mean recombination rate in each of the 129 colonies for which the queen age was known. We applied an ANOVA model to the five age groups (queen age of 0 - 4 years; Figure 2D). As the age groups were unbalanced, we evaluated the significance through permutation tests. We created a null distribution by applying the same ANOVA model to 100 permutations of the age labels and comparing the resulting *F*-statistics to the *F*-statistic from the observed dataset. The same strategy was used in order to evaluate the effect on the recombination rate variance. Both analyses resulted in high *p*-values (0.42 and 0.54, respectively), showing that the maternal age neither has a significant effect on the mean nor the variance of the recombination rate per colony. We also tested for a correlation between the queen age and the mean and variance of recombination rate using Spearman correlation, but the effects were again non-significant.

### GWAS reveals association to mismatch repair gene mlh1

We performed a GWAS analysis by inferring the genotype of each queen and its mean CO rate, based on all sequenced drone progeny. For this analysis we used a more complete set of 3,217,210 SNPs (see Methods). The resulting distribution of *p*-values clearly deviates from the expected distribution as shown in the QQ-plot (Supplementary Figure S4). We corrected for multiple hypothesis testing based on the false discovery rate (FDR) using the method described in (Stephens 2016). At a significance level of *q*-value < 0.05, we found 21 SNPs associated with CO frequency (Figure 5A; Table 2).

**Figure 5.**
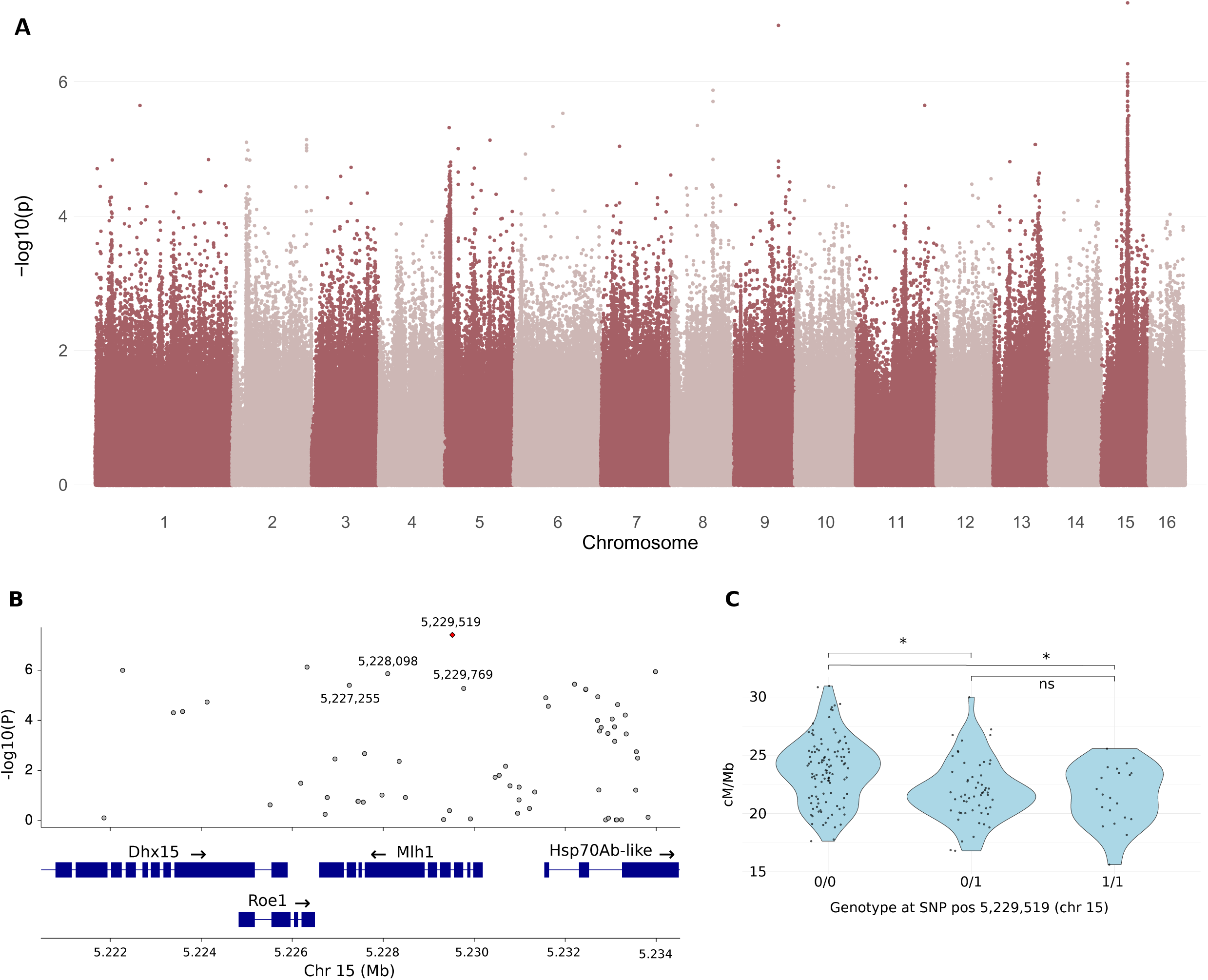
GWAS results for recombination rate. **A)** Manhattan plot of LRT *p*-values from GWAS of recombination rate per colony. **B)** LRT *p*-values of SNPs in the region around *mlh1* (chromosome 15). The four SNPs with the lowest *p*-values within *mlh1* are labeled with their positions and the most significant SNP within *mlh1* is marked in red. **C)** Violin plot showing the recombination rate of each queen, grouped by genotype at the SNP with lowest LRT *p*-value from the GWAS (pos 5,229,519 on chr 15; homozygous reference 0/0, heterozygous 0/1 and homozygous alternate 1/1).

**Table 2.**
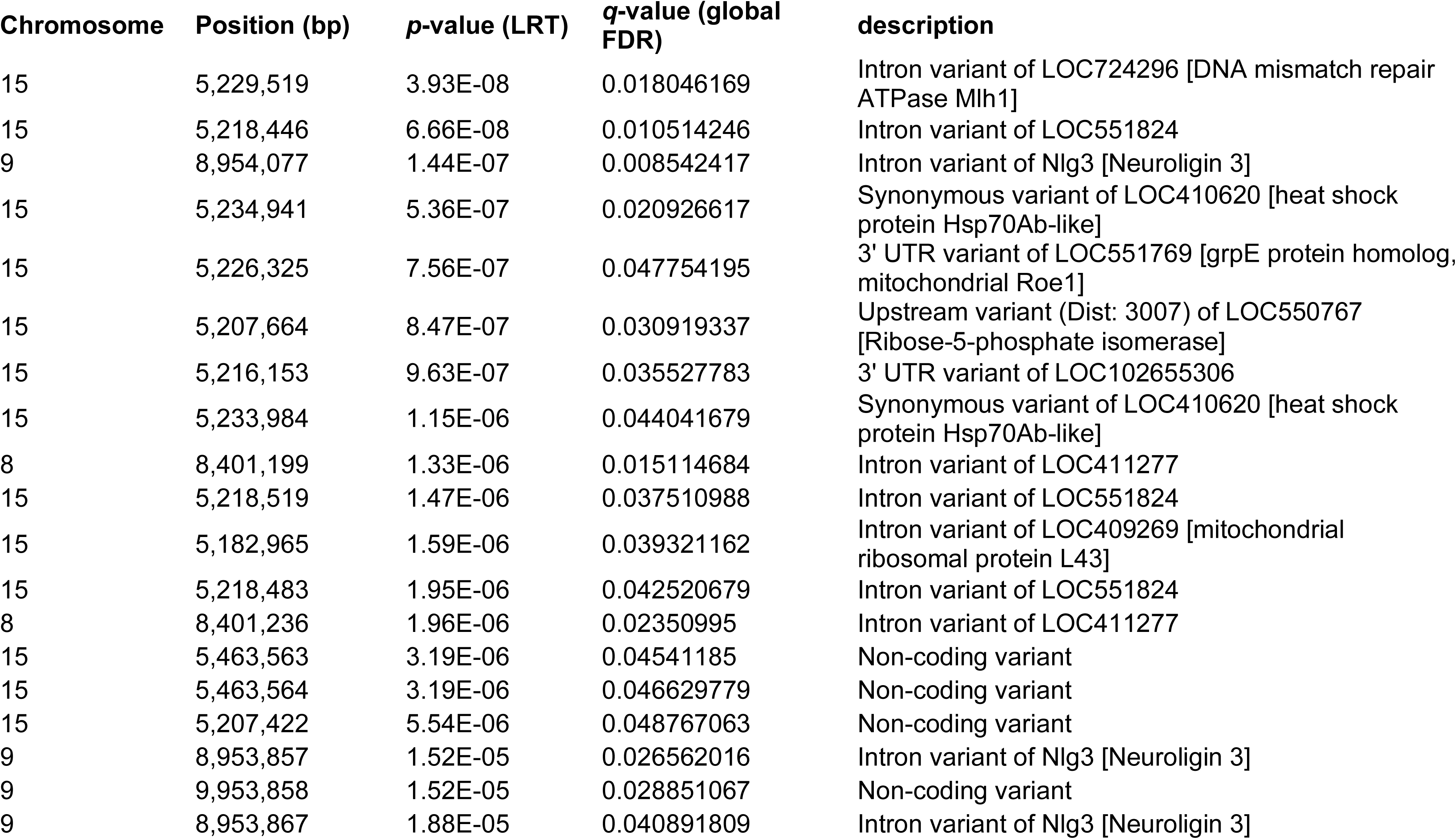
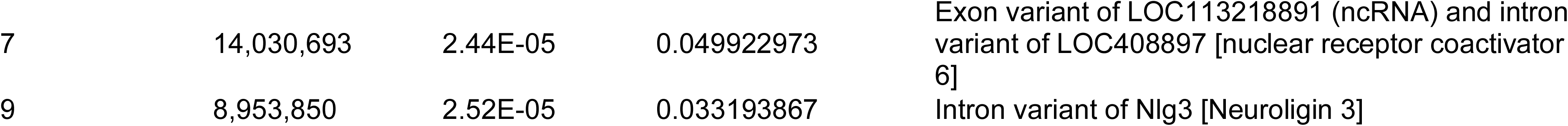
SNPs associated with recombination rate with *q*-value (FDR-adjusted *p*-value) < 0.05 from the GWAS.

The strongest association was found on chromosome 15 around 5,180 - 5,235 kbp, including the gene *mlh1* (5,226,594 - 5,230,184 bp). The SNP with the lowest raw *p*-value (5,229,519 bp; FDR *q*-value 0.018; raw *p*-value 3.93 x 10^-8^) is located within an intron of *mlh1* (Figure 5B). Mlh1 (MutL protein homolog 1) is a core mismatch repair protein across multiple taxa (Baker et al. 1996; Li 2008; Vimal et al. 2018). In yeast, mice, and a wide range of other taxa, Mlh1 forms a dimer with Mlh3, which localises to COs and is required for the resolution of Holliday junctions. We made pairwise comparisons of the average CO rate between the different genotypes at position 5,229,519 and found significant differences between the homozygous reference and the two other genotypes (Figure 5C). The difference between the two homozygous genotypes is 2.4 cM/Mb (10.4%).

There are no exonic missense variants significantly associated with CO frequency in any gene, but within *mlh1* there is one exonic missense SNP which is close to significance (5,228,098 bp; amino acid change D540N; FDR *q*-value = 0.058; raw *p*-value = 1.38 x 10^-6^). A list of all SNPs within *mlh1*, with associated *p*-values, allele frequencies and genetic effects, can be found in Supplementary Table S5. Other genes with significant SNPs in the same peak on chromosome 15 (FDR *q*-value < 0.05) encode heat shock protein hsp70Ab-like (two synonymous variants), mitochondrial grpE protein homolog roe1 (one variant in 3’ UTR), mitochondrial ribosomal protein L43 (one intronic variant) and a few uncharacterized genes (Table 2). There is also one gene on chromosome 9, encoding Neuroligin 3, which has four significant intronic variants.

The Mlh1 protein has two conserved regions separated by a variable region, as identified by multiple sequence alignment of 35 insect species with the NCBI tool COBALT (Papadopoulos and Agarwala 2007), Supplementary Figure S5A; Supplementary Table S6. In a folding prediction from AlphaFold 2 (Jumper et al. 2021), the conserved regions form alpha helices and beta strands while the variable region is an unstructured linker region, Supplementary Figure S5B. The exonic missense mutation with the lowest *p*-value, nucleotide position 5,228,098, D540N, is located within the linker region and is therefore unlikely to have a significant effect on the protein structure or function. There are two missense mutations in the N-terminal conserved region, at nucleotide positions 5,229,327 (V178I) and 5,229,589 (T115I). The former however has a high *p*-value (raw *p*-value = 0.9) and the latter has a very low minor allele frequency (1%) which makes it unreliable for association analyses (Supplementary Table S5). We found no missense variants with strong associations within the conserved domains. A plausible hypothesis is that the causative SNP lies in an intronic or noncoding region and influences the expression of the Mlh1 protein.

## Discussion

Here we estimated intra-specific variation in rates of meiotic recombination and analysed its causes in the honeybee *Apis mellifera*. We analysed whole-genome sequencing data from a set of 1509 haploid drone offspring of 184 females. Using haplotype inference we could estimate the genotypes of the queens and map the location of meiotic crossover events in the genomes of each drone. Our main findings are: 1) We estimated the average rate of crossing over in honeybees to be 23 cM/Mb, with a range of 11 - 41 cM/Mb among haploid males in our dataset. 2) We found substantial variation in rates of recombination among queens compared to among offspring, indicating a narrow-sense heritability of 28%. 3) We found no effect of queen age or genetic background on the crossover frequency. 4) A genome-wide association study for mean crossover rate per queen identified an association with the gene *mlh1*, which forms a known component of the recombination pathway. To our knowledge this is the first locus found to be associated with genome-wide recombination rate variation in an insect.

Strict filtering of the data is necessary for accurate recombination inference, as false variants could lead to inference of false recombination events. We used a relatively sparse set of 326,130 common, high confidence SNPs for this purpose. This is enough for identifying COs, as the number of SNPs greatly exceeds the number of COs on each chromosome (usually < 10) and multiple high-confidence SNPs are expected between each pair of COs. The variation in recombination rate observed between colonies could have both genetic and non-genetic explanations and factors of both kinds have been evaluated in this study. We estimated the broad-sense heritability, i.e. the variation explained by the combined effect of any genetic factors, to 47-52% in the honeybees, which is similar to a previous estimate of 44% in honeybees (Kawakami et al. 2019). Our estimate of the narrow-sense heritability (28%), which specifically includes only the effect of additive genetic factors and controls for random environmental factors, is lower but not close to zero. Our estimates could potentially be biased if there are environmental effects correlated with genetic relatedness that influence recombination rate. However, all sampled colonies experienced similar environmental conditions and the presence of such effects is highly unlikely.

Estimates of the broad-sense heritability in other taxa include 30% in humans (Kong et al. 2004) and 10-40% in different parts of the genome in *D. melanogaster* (Hunter, Huang, et al. 2016). The narrow-sense heritability has been estimated to 46% in mice (Dumont et al. 2009) and lower but significant values in most other studied species (4-13% in cattle (Kadri et al. 2016; Brekke, Johnston, Gjuvsland, et al. 2023); 5-7% in pigs (Johnsson et al. 2021); 12-16% in Soay sheep (Johnston et al. 2016); 12% in female (but insignificant in male) red deer (Johnston et al. 2018); 10-11% in Atlantic salmon (Brekke, Johnston, Knutsen, et al. 2023); and 23% in females and 11% in males of house sparrow (McAuley et al. 2024)). Genetics therefore play an important role in the regulation of recombination rate in most species investigated, in concordance with substantial levels of heritability observed in honeybees. The intra-chromosomal allelic shuffling, although highly associated with CO frequencies, shows much lower heritability, suggesting that the number and distribution of COs along the chromosomes is under genetic control independently of their effects on shuffling genetic variation.

In a previous study, significant differences in recombination rate were found between three populations of honeybees, one European population and two populations from South Africa (subspecies *A. m. scutellata* and *A. m. capensis*), with 28% of the total variation explained by the population differences (Kawakami et al. 2019). The honeybees in our study all had European ancestries, with varying contributions from the C- and M-lineages, but we found no significant difference in recombination rate related to the genomic background, as represented by the first principal component of a PCA. Hence, even though the recombination rate differs between European and African populations and also between African subspecies, the recombination rate in European honeybees does not seem to depend on relative contributions from different European lineages. The role of variation in *mlh1* in determining differences between subspecies is not known, but it is possible that it is an important factor.

The maternal age has been shown to have an effect on the recombination rate in multiple species. In humans, the maternal age and recombination rate are weakly but positively correlated, with increasing age leading to relaxed regulation of the CO events and a lower effect of CO interference (Campbell et al. 2015; Martin et al. 2015). Similar effects have been shown for swine, where the recombination rate increased with increasing maternal parity (Lozada-Soto et al. 2021) and in *D. melanogaster*, where the recombination rate increased with maternal age (Hunter, Robinson, et al. 2016). In mice, the opposite effect has been shown, as increased maternal age led to a decreased number of chiasmata (Polani and Jagiello 1976). A third type of effect has been observed in cattle, where the recombination rate first decreased with maternal age up to 65 months, whereafter it increased instead (Shen et al. 2021). The reason for these varying effects and the prevalence of each type of effect is not well understood. An important difference between mammals and insects is that in female mammals, meiotic recombination occurs already in the foetal stage, when primary oocytes are formed before meiotic arrest. In insects however, oogenesis and meiotic recombination occur throughout life. In this study, we estimated the effect of maternal age based on the 129 queens for which the age was recorded, but we were not able to see any effect on recombination rate in either direction, despite good representation of samples 0-3 years old. This suggests that the recombination rate in honeybees is not strongly influenced by age.

Genome wide association studies have now been used in many species to uncover genetic variation underlying variation in recombination rate. So far, such studies have mainly been applied to domestic and wild animals, with large pedigrees required for estimation of individual recombination rates. Although the genetic basis of crossover variation is likely to be polygenic, several studies, mainly in mammals, have identified loci with major effects. All of these studies have identified hits involved in key meiotic processes and many of those hits are shared between species. The gene *Rnf212* and its paralog *Rnf212B* are involved in the CO/NCO-decision in mice (Reynolds et al. 2013) and have been found significantly associated with recombination rate in humans, cattle, pigs, sheep, Soay sheep, red deer and chicken (Kong et al. 2014; Johnston et al. 2016; Kadri et al. 2016; Petit et al. 2017; Johnston et al. 2018; Weng et al. 2019; Johnsson et al. 2021; Brekke et al. 2022). The MSH4 and MSH5 proteins form a heterodimer which, according to an *in vitro* study of the human homologs, recognizes Holliday junctions and stabilizes the associated recombination intermediates in meiosis I (Snowden et al. 2004). Associations of *MSH4* to recombination rate have been found in humans, cattle and pigs (Kong et al. 2014; Ma et al. 2015; Kadri et al. 2016; Johnsson et al. 2021; Brekke et al. 2022).

The gene *HFM1*, which is associated with recombination rate in cattle and pigs (Kadri et al. 2016; Johnsson et al. 2021), is involved in meiotic CO-formation and synapsis in mice (Guiraldelli et al. 2013). The gene *REC8*, which is one of the main associated genes in Soay sheep and red deer (Johnston et al. 2016; Johnston et al. 2018), controls how chromosomes are paired in synapsis, with deletion of *REC8* leading to synapsis between sister chromatids in mice (Xu et al. 2005). The genes *Rnf212*, *Hfm1* and *Rec8* have also been identified related to recombination rate in mice, however not through GWAS but through QTL-mapping with Haley-Knott regression (Wang and Payseur 2017).

Variation in *mlh1*, the main candidate gene identified in our study, has not previously been implicated in recombination rate variation in any other GWAS, but two different studies in cattle found recombination rate associations with its mammalian interaction partner *MLH3* (Kadri et al. 2016; Brekke, Johnston, Gjuvsland, et al. 2023). The gene *mlh1* is one of multiple eukaryote homologs of the bacterial gene *MutL*, which is involved in mismatch repair (MMR) in *Escherichia coli* (Rayssiguier et al. 1989; Štambuk and Radman 1998). A similar function of *mlh1* in MMR has been shown in yeast (Sugawara et al. 2004; Spies and Fishel 2015) and mammals (Kadyrov et al. 2006; Li 2008). An important role of the MMR machinery is the prevention of recombination between divergent sequences, reducing the risk of inaccurate repair of DSBs, aberrant recombination and chromosomal rearrangements, in a wide range of taxa including bacteria, yeast, mammals and insects (Rayssiguier et al. 1989; LaRocque and Jasin 2010; Do and LaRocque 2015; Spies and Fishel 2015).

In mice, Mlh1 was found to localize to chromosomal CO-sites during meiosis and a null mutation of *mlh1* led to infertility in both males and females (Baker et al. 1996). In yeast (Wang et al. 1999; Zakharyevich et al. 2012) and mammals (Lipkin et al. 2002; Li 2008), Mlh1 forms a heterodimer with Mlh3, called MutLƔ, which is involved in the resolution of Holliday junctions as COs during meiosis. In yeast, various mutations of *mlh3* all resulted in increased numbers of NCOs and one of those mutations disrupted the formation of COs (Al-Sweel et al. 2017). The gene *mlh3* has however been lost from Hymenoptera, Diptera and multiple other insect taxa (Schurko et al. 2010; Tvedte et al. 2017). In *D. melanogaster*, a different protein complex has been shown to have the corresponding function of MutLƔ (Holsclaw et al. 2016; Sekelsky 2017). Depletion of Mlh1 in a hypomorphic strain of *D. melanogaster* caused reduced female fertility, increased number of meiotic DSBs, less efficient repair of DSBs and premature disassembly of the synaptonemal complex (Vimal et al. 2018). Hence, even though the exact function of *mlh1* in honeybees is unknown, its conserved functions in mismatch repair and resolution of Holliday junctions as COs in multiple taxa suggest it likely has similar roles in honeybees.

In mammals, the recombination pathway is relatively well described in terms of the genes involved in different steps, which involves formation and processing of DSBs, strand invasion, synapsis, CO/NCO decision and resolution. The CO frequency could potentially be influenced by the number of DSBs formed, or by variability in other steps along the pathway that influence the probability that a DSB is resolved as a CO (Dapper and Payseur 2019). Intriguingly, in mice, yeast and nematodes, the number of DSBs does not have a major effect on the CO frequency, where there seem to be mechanisms for homeostatic control, which keep the CO frequency relatively unaffected by varying DSB counts (Martini et al. 2006; Chen et al. 2008; Youds et al. 2010; Cole et al. 2012). This suggests that the processes related to the CO/NCO decision are more likely to have a greater influence on recombination rate in these taxa (Dapper and Payseur 2019). Variation in multiple genes related to the CO decision and resolution has been found to affect the CO frequency in mammals (Dapper and Payseur 2019; Johnston 2024). It is therefore likely that variation in *mlh1* in honeybees exerts its effect through this process.

We estimated the genome-wide recombination rate to 23 cM/Mb, which is similar to previous estimates for honeybees, including 26 cM/Mb (Wallberg et al. 2015; Kawakami et al. 2019), 24 cM/Mb (Jones et al. 2019), 18-22 cM/Mb (Fouks et al. 2025), 23 cM/Mb (Solignac et al. 2004), 19 cM/Mb (Hunt and Page 1995; Beye et al. 2006) and 37 cM/Mb (Liu et al. 2015). Other social insects from the order Hymenoptera have similar rates, for example the bees *A. dorsata* (25 cM/Mb; (Rueppell et al. 2016)) and *A. cerana* (17 cM/Mb; (Shi et al. 2013)), the bumblebee *Bombus terrestris* (9 cM/Mb; (Kawakami et al. 2019)) and the ant *Pogonomyrmex rugosus* (11 cM/Mb; (Sirviö et al. 2011)). All those rates are extremely elevated compared with most other metazoan species (Wilfert et al. 2007; Stapley et al. 2017) and compared with non-social insects from the same order, e.g. the solitary bee *Megachile rotundata* (1 cM/Mb; (Jones et al. 2019)) and the solitary wasp *Nasonia* (1.5 cM/Mb; (Niehuis et al. 2010)). This phenomenon has led to multiple hypotheses relating elevated recombination rates to social behaviour, including theories suggesting that recombination, by increasing the genetic diversity, would lead to increased resistance to parasites (Fischer and Schmid-Hempel 2005), facilitate the specialisation on different tasks in the colony (Kent and Zayed 2013) or reduce kin conflicts (Sherman 1979; Templeton 1979). However, a recent study has shown that social behaviour is not always associated with high recombination rates, as termites, which are social insects from a different order, have low recombination rates (around 1 cM/Mb; (Everitt et al. 2025)).

The high recombination rates observed in social hymenopteran species can therefore not be explained by sociality alone. An important step is to investigate how recombination rate is controlled genetically and to identify the loci involved. Our finding that *mlh1* is associated with recombination rate in honeybees raises the question whether this gene could also be related to the differences in recombination rate observed between taxa and if evolution of this gene in social hymenopteran insects might be related to their elevated recombination rates. Further studies comparing *mlh1* between taxa and evaluating evidence of selection are required to answer this question.

## Materials and Methods

### Sample collection

Honey bee drones were collected during the spring and summer of 2019 from multiple locations in Sweden as specified in Supplementary Table 1. Honeybee colonies that were known to be closely related were not included in the sampling. To collect drones, we first cut a section of wax containing drone brood (larvae or pupae) from each colony. Each piece of brood was then placed in an incubator at 35 °C in a separate cardboard box until adult bees emerged, which were used for DNA extraction.

### DNA extraction and library preparation

Tissue samples were collected from the thorax of the drones. The exoskeleton was removed using a sterile scalpel in a petri dish placed on a cold surface. The tissue was immediately transferred into an ATL buffer (Qiagen) for lysis according to Qiagen DNeasy Blood and Tissue Kit protocol with individual spin columns. The standard protocol was adapted by including an extended lysing time and a RNase step. For a subset of the samples, alternative protocols were used: the Qiagen DNeasy Blood and Tissue 96-well format (13 drones, 2 colonies) and an isopropanol-based extraction method (52 drones, 7 colonies; Supplementary Table S1). Library preparation was done with the Illumina Nextera Flex kit following standard protocols.

### Sequencing

Sequencing with 2 x 150 bp paired end reads was performed on a NovaSeq 6000 instrument S4 flow cell for most samples and on a HiSeqX instrument using HiSeqX SBS chemistry for 112 drones and 15 colonies (Supplementary Table S1).

### Mapping, variant calling and filtering

The 1561 sequenced drones and previously published sequences from 158 additional drones (PRJNA516678) (Kawakami et al. 2019) were included in the initial dataset. Samples from two reference populations (nine *A. mellifera carnica* worker bees from Eastern Europe (Harpur et al. 2014) and 17 *A. mellifera iberiensis* drones from Iberia (Henriques et al. 2018)) were also included for comparison in the PCA analysis and were processed together with the main drone samples during the first steps. All samples were mapped to the Amel_HAv3.1 reference genome (Wallberg et al. 2019) using the Burrows-Wheeler alignment tool BWA version 0.7.18 (Li and Durbin 2009) with the BWA-MEM algorithm. The resulting BAM-files were sorted and indexed with SAMtools package version 1.20 (Li et al. 2009). Read groups were added and duplicate reads were removed using Picard Toolkit version 3.1.1 (Broad Institute 2019) with the tools AddOrReplaceReadGroups and MarkDuplicates, respectively. Variants were called using GATK version 4.6.1.0 (Poplin et al. 2017; Auwera and O’Connor 2020), following their best practices workflow for germline short variant discovery (https://gatk.broadinstitute.org/hc/en-us/articles/360035535932-Germline-short-variant-discovery-SNPs-Indels, last accessed 2025-06-10) with the tools HaplotypeCaller, GenomicsDBImport and GenotypeGVCFs, leading to a set of 13,164,096 variants.

We treated all sequences as diploid during variant calling even though most of them are actually haploid, in order to identify low quality sites that are called as heterozygous in the haploid drones. As one of the reference populations contains diploid individuals, those were treated differently when filtering for heterozygosity, as described in the section about PCA.

The main dataset without the reference populations was filtered as follows. Only bi-allelic SNPs and no indels were included and those were first subject to hard filters applied with the GATK tool VariantFiltration. The hard filtering limits applied were all equal to or stricter than the recommended limits and adapted to the distributions of the statistics (QD<20, FS>60, MQ<40, MQRankSum<(−12.5), ReadPosRankSum<(−8), SOR>3). In addition, sites with a total depth (INFO/DP-value) ≥ 20,000 were removed, as extremely high read depth is indicative of mapping errors and 99% of the sites have a total depth of <16,000 (mean DP = 12,070). The genotype qualities (GQ-values) were recalculated in order to reflect the fact that there should be no heterozygous genotypes in the data, so that the GQ-value is equal to the difference in likelihood between the two homozygous genotypes. The updated values were then used for filtering on GQ ≥20, changing the genotypes with lower qualities to missing data. Sites with >10 heterozygous genotypes were removed and remaining heterozygous genotypes were encoded as missing data. Finally, sites with >10% missing genotypes were also removed, leaving a set of 326,130 high quality SNPs.

Out of the 1719 drone samples, 137 individuals were excluded due to various quality issues: high heterozygosity (>3% of SNPs heterozygous), missing data at many sites (individuals with missing genotypes at >95% of the SNPs and colonies where less than five individuals had >50% called genotypes), low relatedness to the rest of the colony (as explained below) and colonies where less than five individuals remained, as the recombination inference is less reliable when there are fewer individuals to base the inference on. We also removed 73 drones that were inferred to have an extreme number (>100) COs based on the YAPP analysis (see below). These were considered to be false positives as 95% of the drones have less than 75 COs in total. The final dataset therefore consisted of 1509 drones from 184 different colonies.

The relatedness between the samples was evaluated within and between colonies, in order to identify drones not belonging to the expected colony. This was done using the VCFtools function --relatedness, which estimates the parameter *A_jk_* as defined by (Yang et al. 2010). In order to limit the memory requirements, the calculations were done separately on four different, approximately equally sized sets of individuals, comparing all pairs of individuals within each set. For each individual, the mean relatedness to its siblings from the same colony was compared to the maximum relatedness to any other individual in the same set. Misassignment was suspected when the latter estimate was higher than the former and led to the exclusion of one colony.

A few colonies and individuals were sequenced twice (17 drones from four different colonies; Supplementary Table S2) in order to evaluate the reliability of the recombination inference with regard to sequencing results. All replicates were confirmed by the relatedness score *A_jk_* to be from the same respective individuals.

### Recombination inference with YAPP

The strictly filtered dataset of 326,130 SNPs was used for recombination inference with the software YAPP version 0.4.2 (Servin 2025) and Python version 3.9. This was done in three steps, with the tools sperm, phase and recomb. All tools were run with an expected recombination rate --rho of 26 cM/Mb, as an average of recent estimates (Wallberg et al. 2015; Jones et al. 2019; Kawakami et al. 2019), and a genotyping error rate --err of 0.01. The tool sperm was run in order to infer the sequence of each queen based on the sequences of the drones from the same colony, as the drones are haploid outcomes of meiosises in the queen and in this way equivalent to sperm cells. The resulting phased queen sequences were merged with the initial VCF-file. The merged VCF-file with both queens and drones was then used as input to the tool phase, which was run in order to create the necessary input files for the final step, i.e. the recombination inference. The recombination events were inferred with the tool recomb, which outputs the chromosomal coordinates of each recombination event in each offspring as well as the region of each chromosome where recombination inference could be made, which is usually not the full chromosome due to the number of markers being limited. The recombination rate per drone was calculated as the number of recombination events divided by the length of the informative portion of the genome. The recombination rate of each queen was calculated as the mean of the recombination rates of the drones in the corresponding colony.

### Recombination rate variation along chromosomes

The output from YAPP specifies an interval in bp for each CO in each drone, where the start and end of each interval corresponds to different SNP-positions, between which the haplotype switches. The total recombination rate between each pair of consecutive SNPs was calculated in R as the sum of the overlapping recombination intervals from all drones, with each overlapping interval adding a value of one divided by the length of the interval. In order to convert those rates to cM/Mb, they have to be divided by the total number of drones (1509).

### Heritability analysis

The heritability of the recombination rate was estimated in two different ways, based on ANOVA and based on restricted maximum likelihood (ReML), respectively.

The broad-sense heritability *H^2^* was calculated based on a one way ANOVA of the recombination rate per drone with the colony information as a random effect factor using R version 4.2.3 (R Core Team 2023) with the function aov (Hastie 2017). The sums of squares for the colony factor and residuals, *SS_colony_* and *SS_residual_*, were used to calculate *H^2^* as specified in Eq. 1 below.

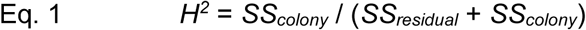

The broad- and narrow-sense heritabilities (*H*^2^ and *h*^2^) were also calculated with the following model (Eq. 2). If the number of drones is *N* and the number of colonies is *Q*, then *Y* is the (*N*\*1) vector of observed recombination rate per drone (scaled to a Gaussian distribution in order to avoid numerical problems), 𝜇 is the intercept, *m* is the (*Q*\*1) vector of environmental effect per colony and *a* is the (*Q*\*1) vector of additive genetic effects per colony. *Z* is the (*N*\**Q*) incidence matrix relating the colony effects to the drones and *e* is the (*N*\*1) vector of residuals.

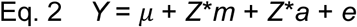

This was modelled in a restricted maximum likelihood (ReML) framework with *m* and *a* as random effect factors using the R package MM4LMM (Johnson and Thompson 1995; Laporte et al. 2022) with the function MMEst. A genomic relationship matrix (GRM) between the queens was included as a covariance matrix for the additive genetic effects. The GRM was computed with LDAK v. 6 (Speed et al. 2017; Speed et al. 2020), using the functions cut-weights and calc-weights-all, which thin the variants to an *r*^2^-threshold of 0.98 and weight them by linkage disequilibrium and minor allele frequency, followed by the function calc-kins-direct with the LDAK-Thin model to calculate the pairwise kinships between the queens. The variances estimated by MMEst (𝜎_m_, 𝜎_a_, 𝜎_e_ for the factors *m*, *a* and *e*) were then used for calculating the heritabilities (Eq. 3-4)

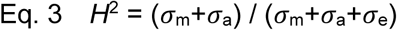

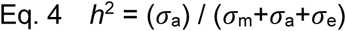

### Analysis of intra-chromosomal allelic shuffling

Intra-chromosomal allelic shuffling, *̄r*, as defined by (Veller et al. 2019), was calculated in R for each drone based on the equation below, where *n* is the total number of chromosomes (*n* = 16 for honeybees), *p*_k_ and (1-*p*_k_) are the proportions of alleles from each parental origin and *L*_k_ is the chromosome length as a fraction of the genome length.

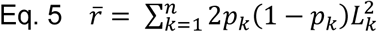

The heritability of the allelic shuffling was calculated within an ReML framework as described above for the recombination rate (Eq. 2) and also with an ReML model controlling for CO frequency. In the latter case, the allelic shuffling phenotype (*Y*) was replaced by the residuals from linear regression of the allelic shuffling on the CO frequency.

### Analysis of crossover interference

Crossover interference was analyzed using the R package xoi version 0.72 (Broman and Weber 2000) and the function fitStahl, which utilizes the Houseworth-Stahl interference escape model (Housworth and Stahl 2003). This model takes into consideration both class I COs (affected by interference) and class II COs (not affected by interference). The strength of CO interference and the proportion of non-interfering COs in each queen was estimated based on the genetic positions of the COs in her drone offspring. For each drone, the genetic distances between the COs were inferred based on the physical positions of the COs obtained from YAPP and the number of intervening COs in the total set of drones.

### PCA

In order to analyse the population structure among the sampled drones, sequence data from two reference populations were included for comparison: nine *A. mellifera carnica* worker bees from Eastern Europe (Harpur et al. 2014) and 17 *A. mellifera iberiensis* drones from Iberia (Henriques et al. 2018), representing the C-group and M-group, respectively, which are the main ancestries expected among the sampled drones (Wallberg et al. 2014). The processing of the reference populations was done mainly as described for the main drone dataset in the section about mapping and variant calling above, with the difference that the heterozygosity filter could not be applied to the diploid worker samples.

The VCF-file of the drones and reference populations was converted to the format required for PCA (.ped and .map files) using PLINK 2 v2.0.0-a6.9LM (Chang et al. 2015; Purcell and Chang). The dataset was additionally filtered for minor allele frequency ≥ 0.1 at each site across all samples, as well as pruned for LD with the PLINK function –indep-pairwise to remove SNPs with *R^2^* >0.2 to any other SNP in a window of 50 SNPs, with a step-size of 5 SNPs, leading to a set of 295,812 SNPs. The eigenvectors and eigenvalues for the PCA were calculated with the PLINK function –pca and then visualized in R.

### Effect of queen age

The association between the mean recombination rate per colony and the age of the queen was estimated in R both with Spearman correlation and one way ANOVA, as we did not know which type of relationship to expect. As the age groups are highly unbalanced with the majority of queens in the lower ages, the results from those tests are difficult to interpret and therefore we also made permutation tests. The queen age labels were randomly shuffled 100 times, calculating Spearman’s *⍴* and the ANOVA *F*-value each time. The p-values were then calculated as the proportion of *⍴* and *F*-values from the randomized tests that were equal to or more extreme than the corresponding observed values. The association between the variance in recombination rate per colony and the queen age was estimated in the same way.

### Reconstruction of queen sequences for GWAS

For the recombination inference, a strictly filtered set of SNPs was used, as incorrectly called SNPs could lead to false positives. For the GWAS analysis, a greater number of SNPs facilitates fine-mapping and identification of putatively causative variants. Therefore the set of SNPs used in this case was filtered more leniently while still assuring high quality. The limits used for this dataset differed from the limits described for the main dataset in the following ways: the hard filters for QD and FS were changed from <20 to <2 and from >60 to >200, respectively, and the filter for missingness per site was changed from ≤10% to ≤30%. This resulted in 3,213,079 biallelic SNPs in the drones and those were used for reconstruction of the diploid queen sequences using a custom script (https://github.com/Jimi92/QueenGT_inference) as described below.

We reconstructed the genotype of each queen using the drone sequences from the corresponding colony, as the drones are mosaics of the queen’s haplotypes. For each SNP, the queen genotype was inferred to be either homozygous for the major allele among the drones or heterozygous if both alleles were present at a certain minimum frequency among the drones.

Due to the limited number of sequenced drones per colony, a challenge of this approach is to define which minor allele frequency is required in the colony for the queen to be considered heterozygous at each position. Most colonies have eight drones, which means that for a locus where the queen is heterozygous, there is a 93% chance that at least two drones in her colony carry each of the two alleles. Therefore we set the minor to major allele ratio to 0.25 meaning that at least two out of eight drones need to have the minor allele for the site to be called as heterozygous in the queen.

### GWAS

GWAS was performed with GEMMA v.0.98.5 (Zhou and Stephens 2012) based on the reconstructed diploid queen sequences (described above), with a relatedness matrix as covariate and the recombination rates estimated by YAPP as phenotypes.

First, the set of 3,213,079 SNPs was filtered using PLINK 2.0 with the following limits: individual missingness ≤10%, genotype missingness ≤10%, Hardy-Weinberg equilibrium exact test *p*-value ≥0.001 and minor allele frequency ≥1%, resulting in a set of 2,762,043 SNPs and 184 queens, which were used as input to GEMMA. After some additional filters applied by GEMMA with the default parameters, except for MAF which was changed to 0.05, 2,310,593 SNPs remained and were used in the analyses. A centered genotype relatedness matrix was calculated with the option ‘-gk 1’ and then included in the association analysis in order to control for population structure. The association analysis was conducted using a univariate linear mixed model (recombination-rate ∼ genotype + (1|relatedness)). The *p*-values from the likelihood ratio test (LRT) were visualized in a Manhattan plot using the R package locuszoomr (Lewis and Wang 2024). In order to account for multiple hypothesis testing, FDR-adjusted *q*-values were calculated with an adaptive shrinkage method using the ashr R package version 2.2-63 (Stephens 2016). SNPs with a *q*-value of less than 0.05 were deemed significant.

### Analysis of candidate SNPs

The highest scoring SNPs were further analyzed with the Ensembl Variant Effect Predictor (VEP) (McLaren et al. 2016), the NCBI *A. mellifera* annotation release 104 accessed through Ensembl Metazoa release 60 (Dyer et al. 2025) and with the UCSC Genome Browser (Perez et al. 2025), in order to identify their locations relative to functional elements and genes and their potential biological effects.

### Multiple sequence alignment and structure prediction of the Mlh1 protein

Mlh1 conservation was investigated by protein-level multiple sequence alignment (MSA) of Mlh1 homologs from 35 different insect species, whose transcript IDs were available on NCBI (Supplementary Table S6). The alignment was done using the NCBI Constraint-based Multiple Alignment Tool (COBALT) (Papadopoulos and Agarwala 2007) with the default parameters.

The structure of the *A. mellifera* Mlh1 protein was predicted using AlphaFold 2 (Jumper et al. 2021). Visualization and labeling of the predicted protein structure was done using the web-based NCBI tool I see in 3D (iCn3D) (Wang et al. 2022).

## Data and Code availability

Scripts and code available on GitHub: https://github.com/turideveritt-code/RecRateGWAS.git. All data have been uploaded to ENA with the ProjectID PRJEB105756.

## Acknowledgements

We acknowledge grant 2018-03896 from The Swedish Research Council to MTW. No funders had any role in study design, data collection and analysis, decision to publish, or preparation of the manuscript. We acknowledge help from several beekeepers in Sweden. The authors acknowledge support from the National Genomics Infrastructure in Stockholm funded by Science for Life Laboratory, the Knut and Alice Wallenberg Foundation and the Swedish Research Council, and NAISS/Uppsala Multidisciplinary Center for Advanced Computational Science for assistance with massively parallel sequencing and access to the UPPMAX computational infrastructure. The computational analysis was enabled by resources provided by the National Academic Infrastructure for Supercomputing in Sweden (NAISS), partially funded by the Swedish Research Council through grant agreement no. 2022-06725.

## Author contributions

M.T.W. designed the project. M.T.W and J.d.M. collected samples. A.O. performed DNA extractions. T.E., L.H.T., D.T., T.R. performed data analysis. T.E. and L.H.T. produced the figures. B.S. contributed and advised analysis methods. T.E. wrote the manuscript with input from M.T.W and all other authors.

## Supplementary Tables

**Supplementary Table S1)** Data per colony: Sampling details, queen age, number of drones in colony, number of informative SNPs (where each allele is present in at least one drone), mean and variance of the CO frequency.

**Supplementary Table S2)** Data per drone: Quality statistics, read depth, number of CO:s, size of the part of the genome where recombination inference could be made, CO frequency.

**Supplementary Table S3)** Relatedness parameter *A_jk_*, which is on average 1 for an individual with itself and 0 for unrelated individuals (Yang et al. 2010): mean relatedness of each individual to its siblings from the same colony and maximum relatedness to any individual from another colony. Colonies are divided into four different sets for the calculations in order to reduce memory requirements.

**Supplementary Table S4)** Total number of CO per bp, summed across all 1509 drones, calculated between each pair of consecutive SNPs.

**Supplementary Table S5)** All SNPs found within the gene *mlh1*.

**Supplementary Table S6)** Table of species and accession numbers of mlh1 homologs used in multiple sequence alignment (Supplementary Figure S5A).

## Supplementary Figures

**Supplementary Figure S1)** Heatmaps of the relatedness score *A_jk_* (Yang et al. 2010) between drones in four different sets, with stronger red color indicating higher relatedness.

**Supplementary Figure S2)** Positions of recombination events along each chromosome in replicate samples. There are two samples from each individual, drawn in the same color. As recombination events are inferred as intervals rather than exact positions, the start of each interval is marked by an asterisk and the end is marked by a diamond.

**Supplementary Figure S3)** Recombination rate per position along all chromosomes based on data in Supplementary Table S4. Pericentromeric regions identified in (Wallberg et al. 2019; Everitt et al. 2023) are marked by dashed red and green lines for start and end of the region, respectively.

**Supplementary Figure S4)** QQ-plot of observed LRT *p*-values from recombination rate GWAS versus expected *p*-values from random distribution.

**Supplementary Figure S5)** Structure and evolution of the *mlh1* gene. **A)** Protein-level multiple sequence alignment of *mlh1* orthologs from 35 different insect species (listed in Supplementary Table S6). Highly conserved positions are shown in red, lower conservation in blue and regions with gaps in the alignment are shown in gray. **B)** Predicted structure of the *A. mellifera* Mlh1 protein from AlphaFold 2. Regions are colored by the confidence of the prediction (pLDDT score). Amino acids affected by missense SNP:s are marked in green and labeled with their positions.

