## Supplementary Figure for "Genetic determinants of intraspecific variation in crossover frequencies in the honeybee, *Apis mellifera*"

**for**

A)

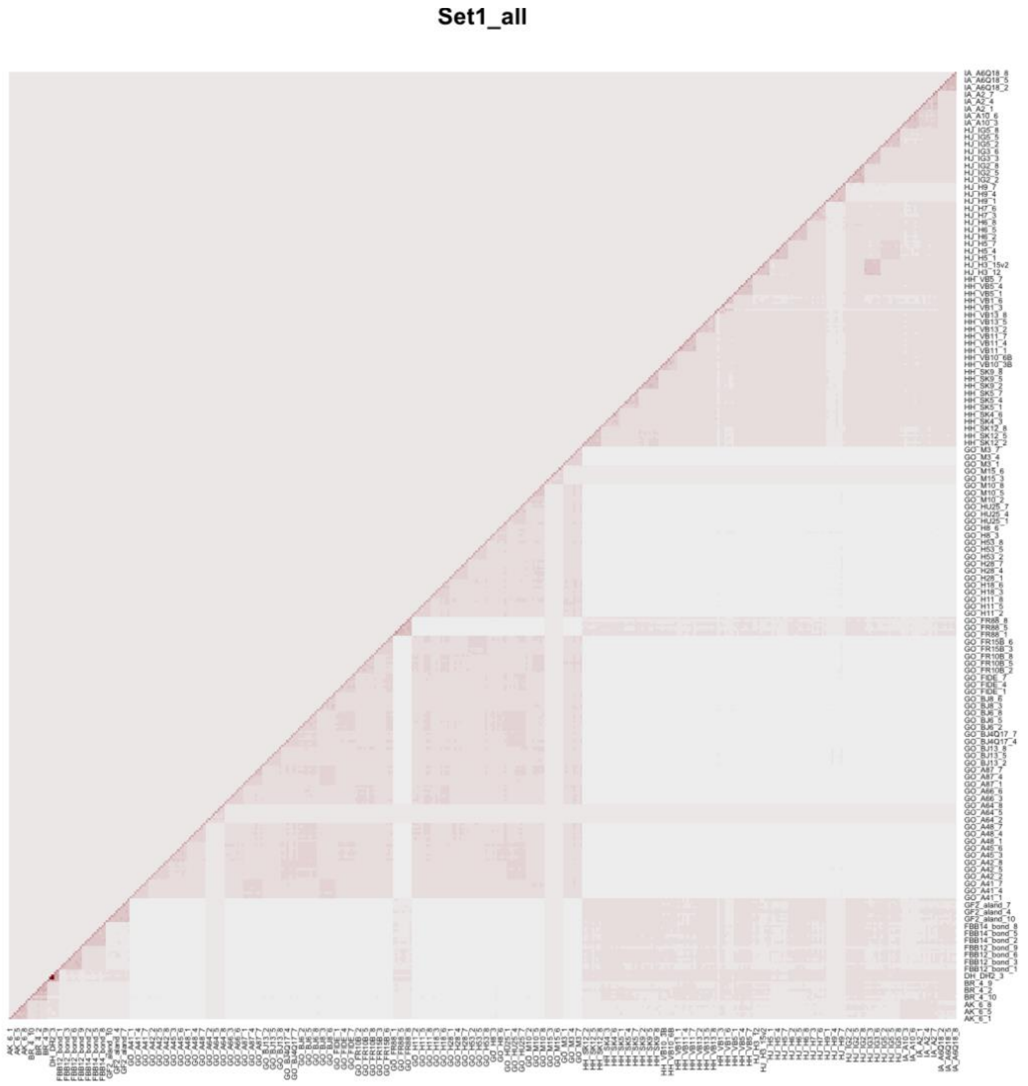

B)

**Set2_all**

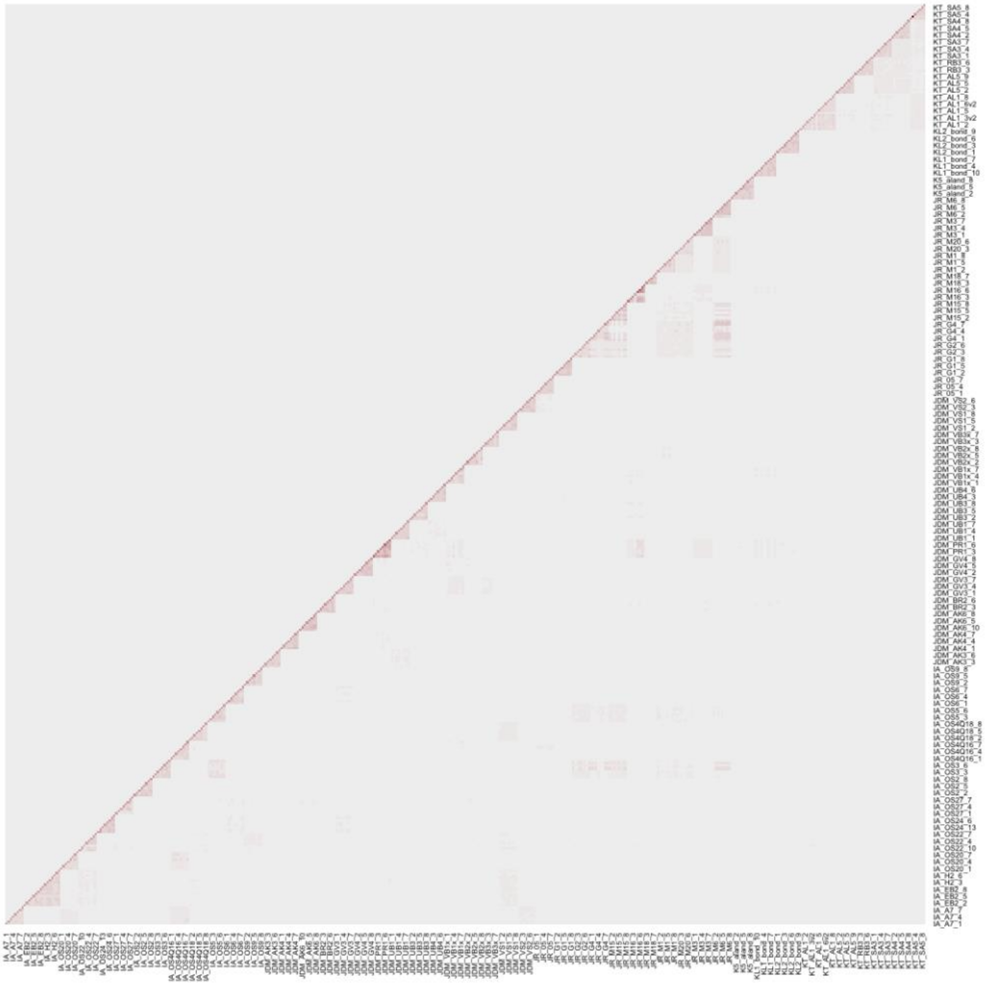

C)

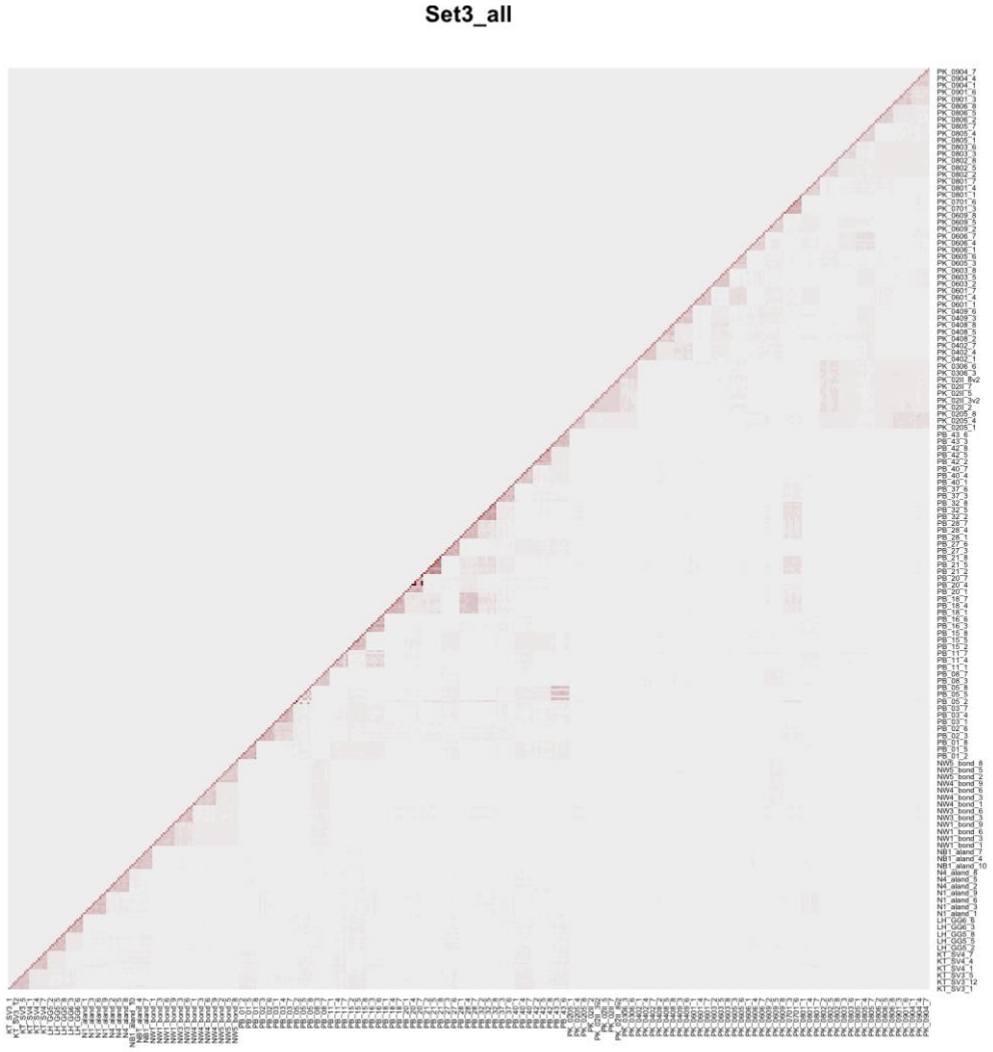

D)

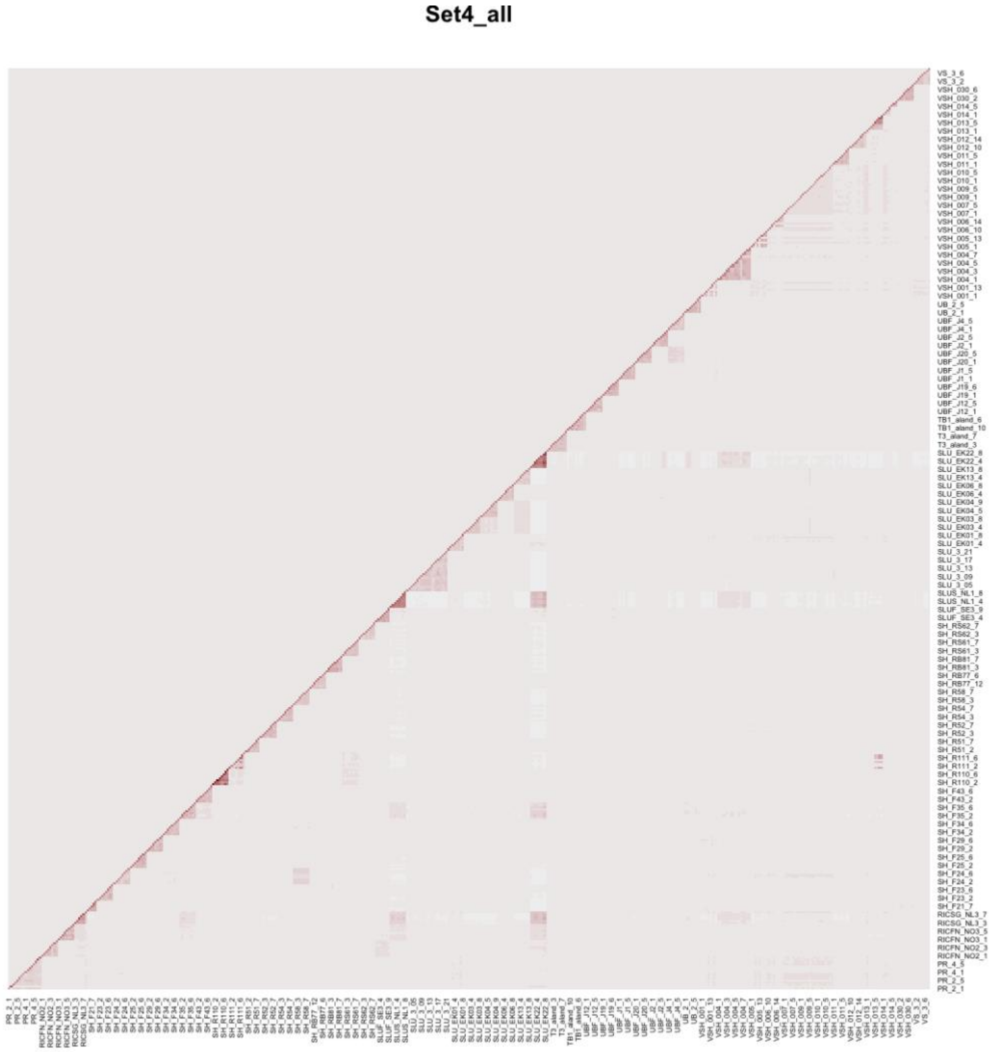


**Supplementary Figure S1)** Heatmaps of the relatedness score $A_{jk}$ (Yang et al. 2010) between drones in four different sets (A-D), with stronger red color indicating higher relatedness.

A)

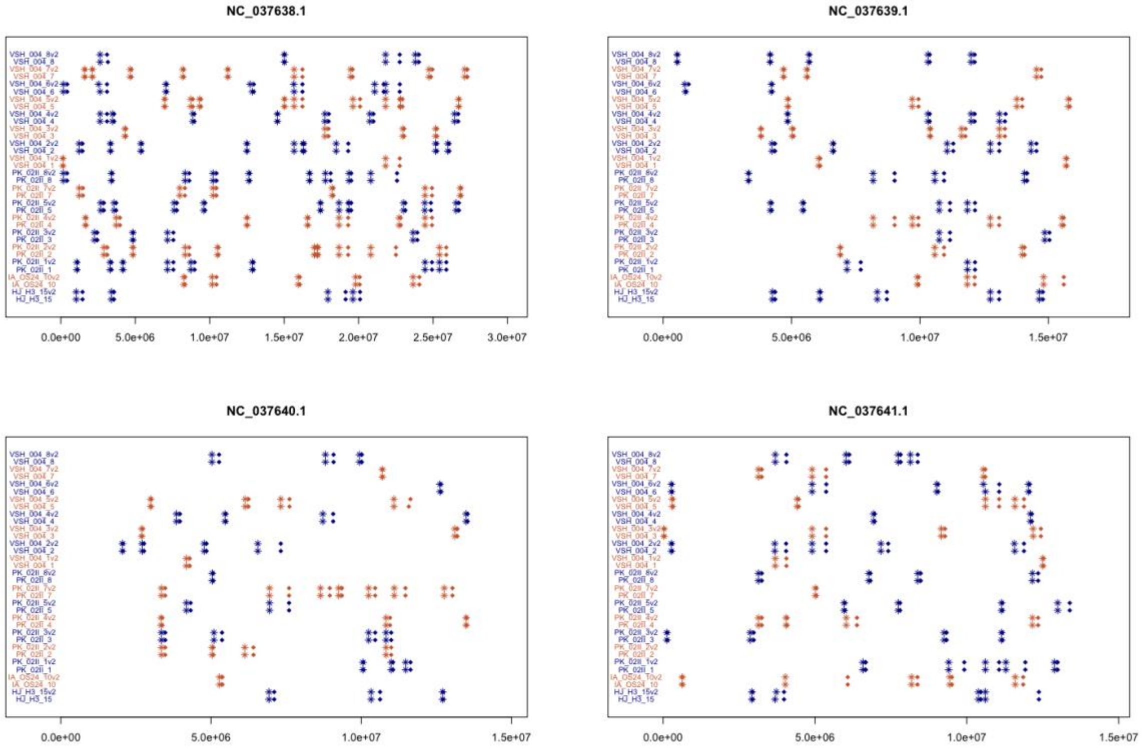
NC_037638.1
0.0e+00
5.0e+06
1.0e+07
1.5e+07
2.0e+07
2.5e+07
3.0e+07
NC_037639.1
0.0e+00
5.0e+06
1.0e+07
1.5e+07
NC_037640.1
0.0e+00
5.0e+06
1.0e+07
1.5e+07
NC_037641.1
0.0e+00
5.0e+06
1.0e+07
1.5e+07

B)

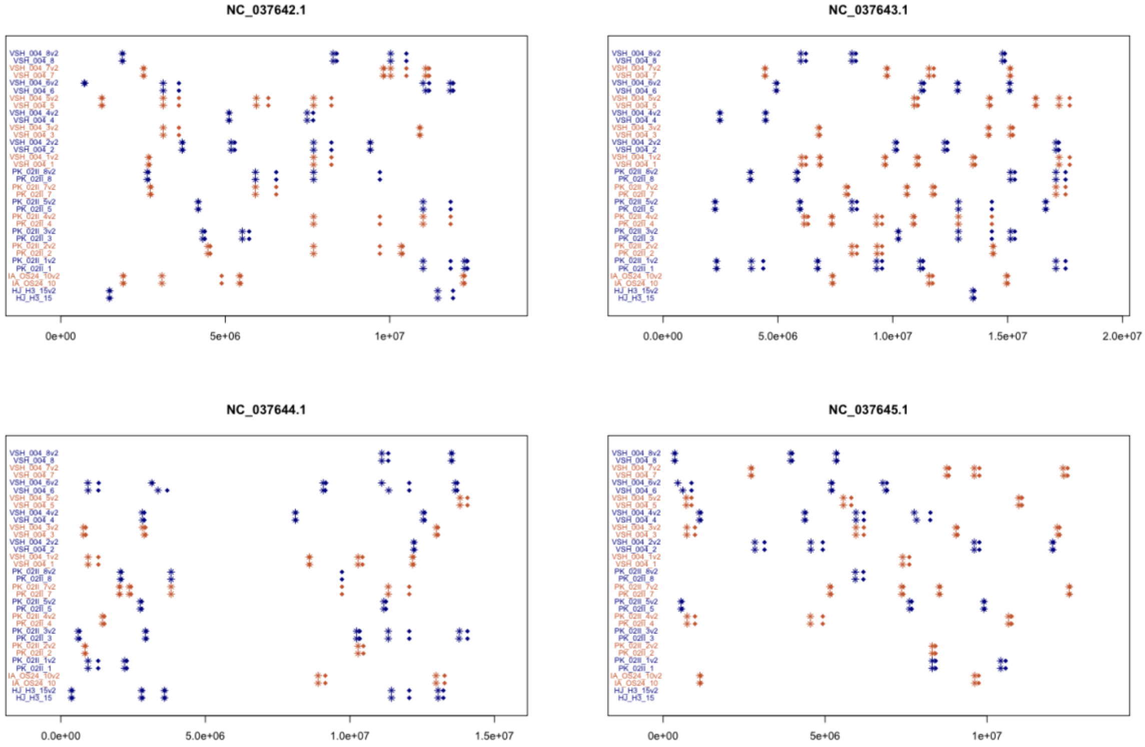
NC_037642.1
0e+00
5e+06
1e+07
NC_037643.1
0.0e+00
5.0e+06
1.0e+07
1.5e+07
2.0e+07
NC_037644.1
0.0e+00
5.0e+06
1.0e+07
1.5e+07
NC_037645.1
0e+00
5e+06
1e+07

C)

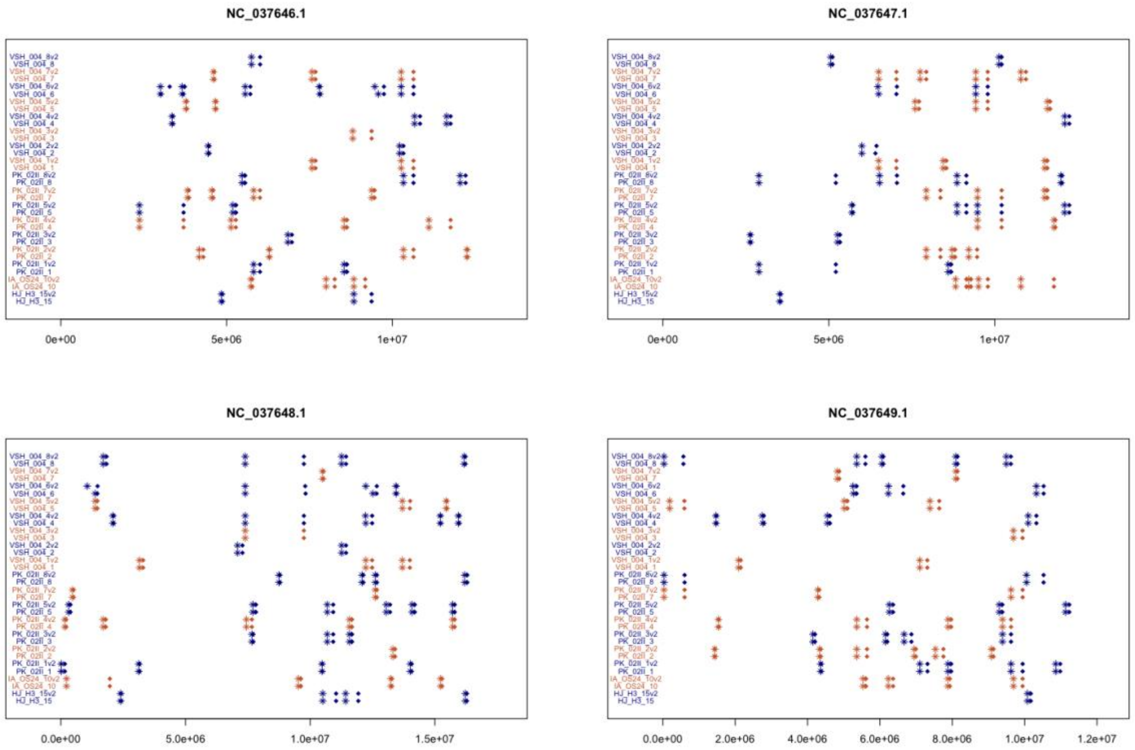


D)

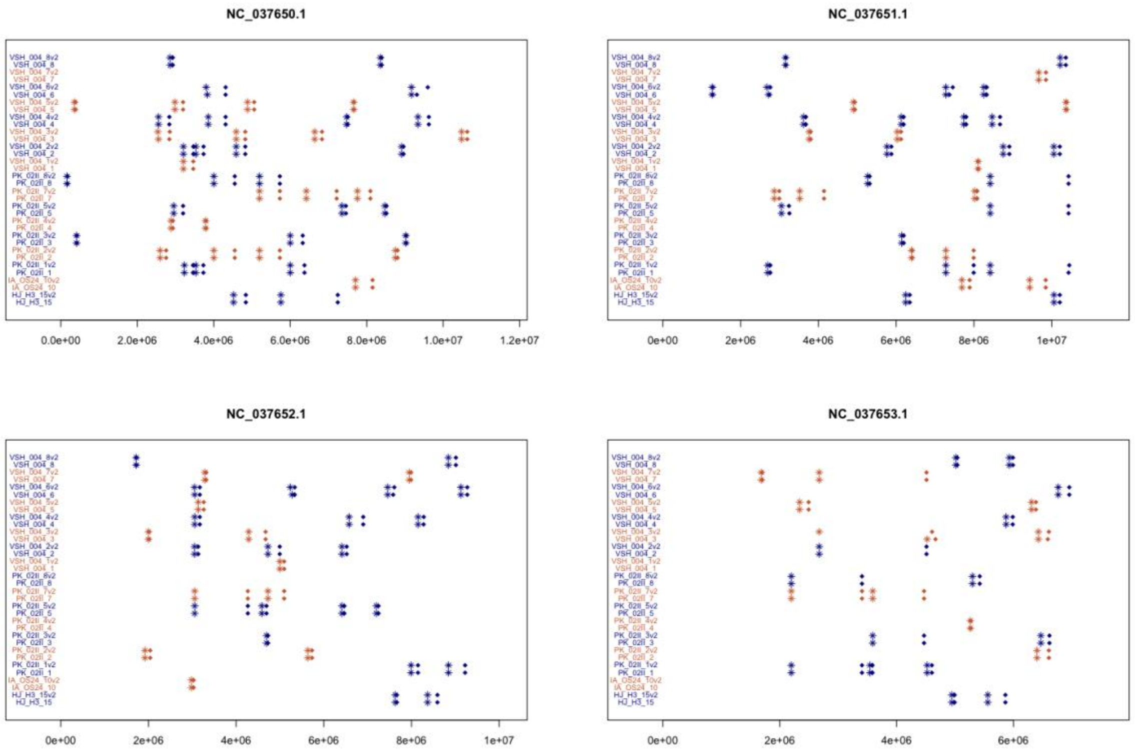


**Supplementary Figure S2)** Positions of recombination events along each chromosome in replicate samples. There are two samples from each individual, drawn in the same color. As recombination events are inferred as intervals rather than exact positions, the start of each interval is marked by an asterisk and the end is marked by a diamond.

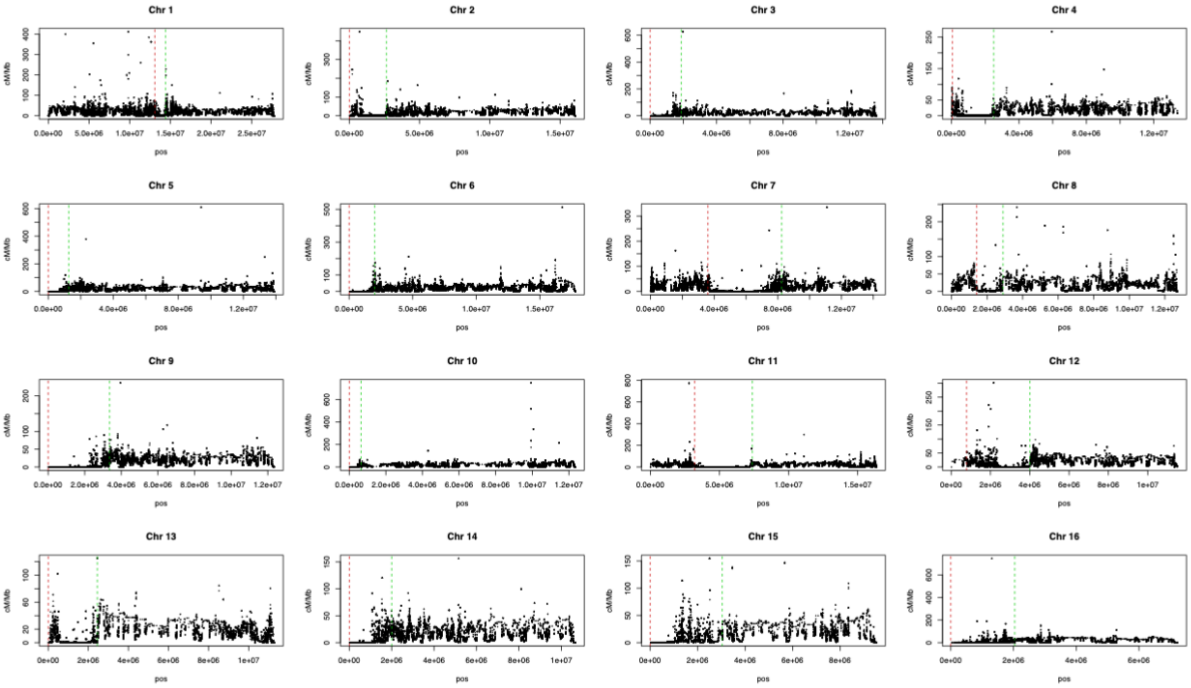


**Supplementary Figure S3)** Recombination rate per position along all chromosomes based on data in Supplementary Table S4. Pericentromeric regions identified in (Wallberg et al. 2019; Everitt et al. 2023) are marked by dashed red and green lines for start and end of the region, respectively.

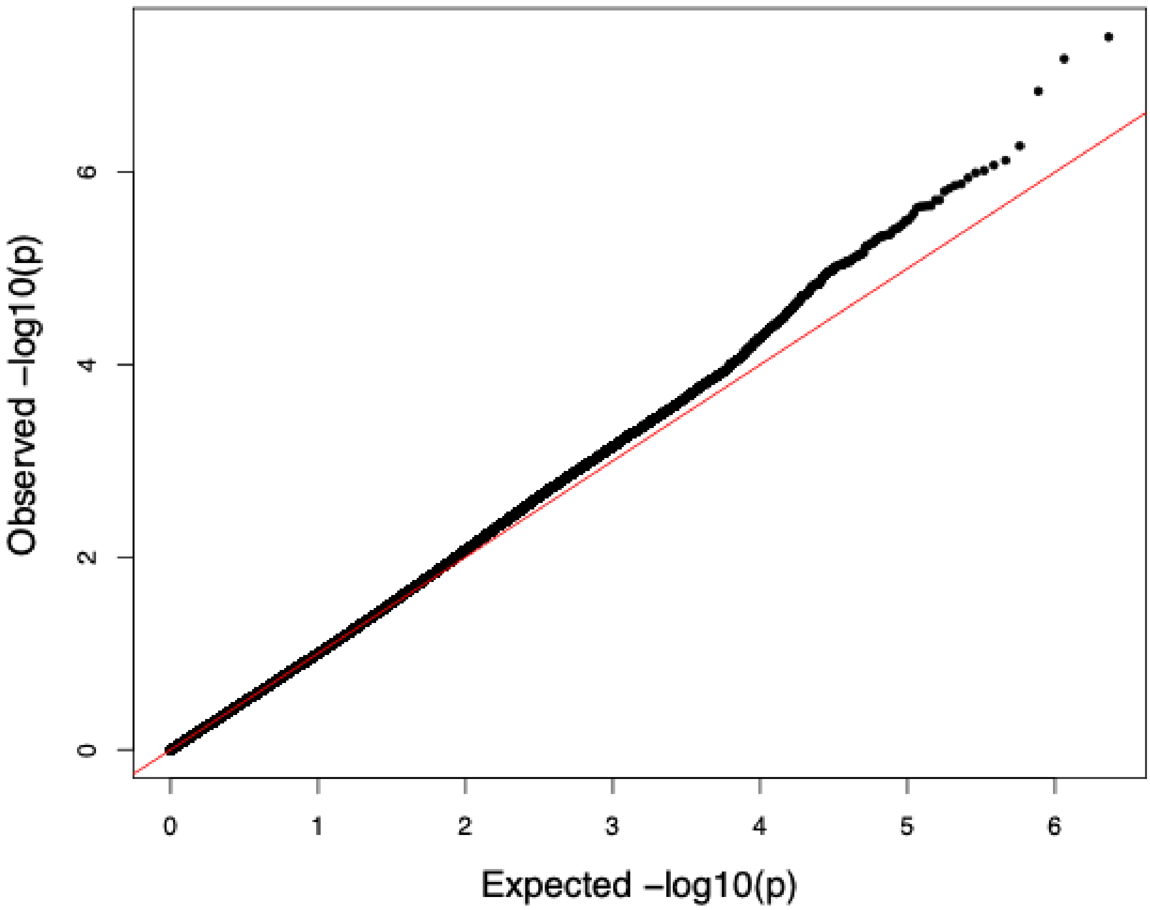


**Supplementary Figure S4)** QQ-plot of observed LRT *p*-values from recombination rate GWAS versus expected *p*-values from random distribution.

**A)**

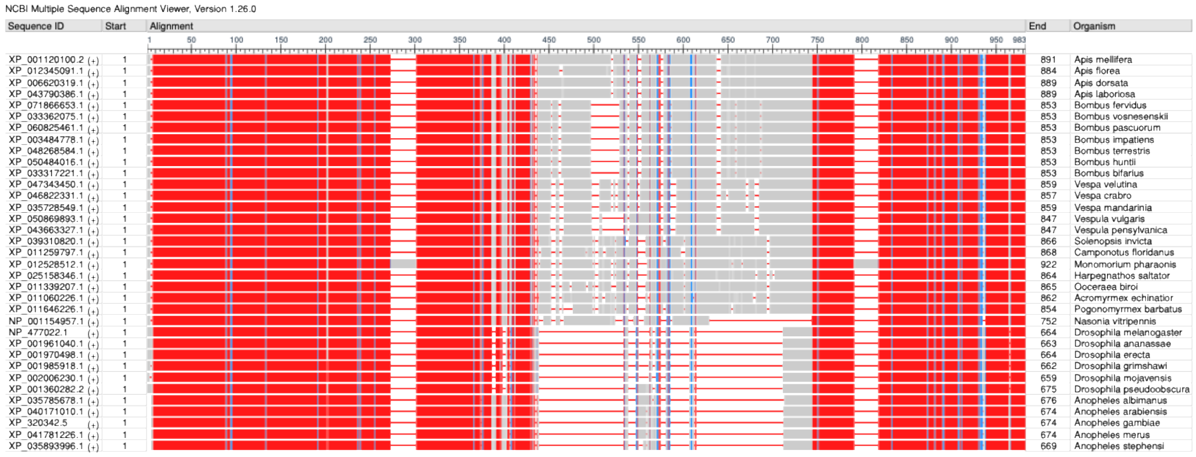


**B)**

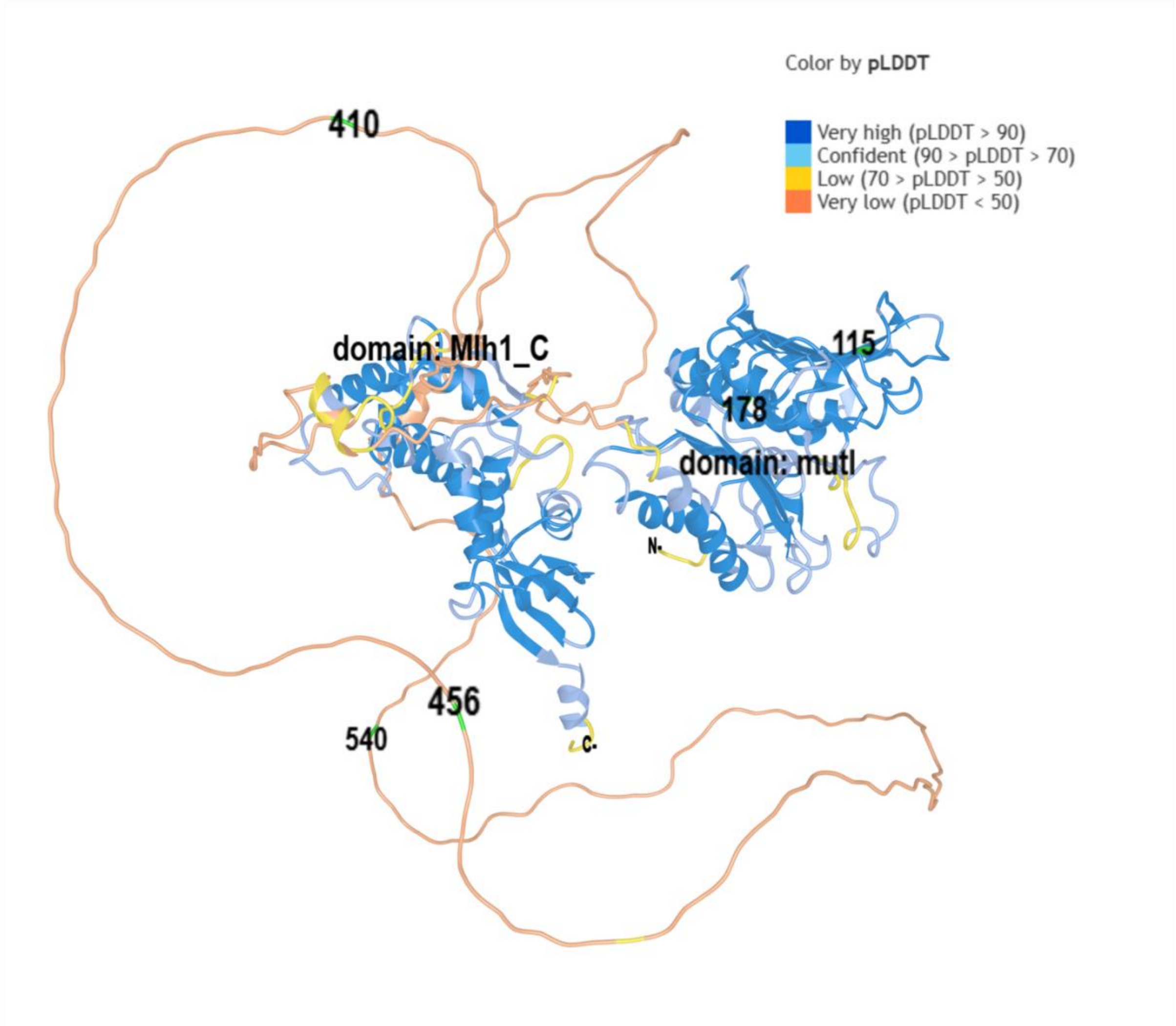


**Supplementary Figure S5)** Structure and evolution of the *mlh1* gene. **A)** Protein-level multiple sequence alignment of *mlh1* orthologs from 35 different insect species (listed in Supplementary Table S6). Highly conserved positions are shown in red, lower conservation in blue and regions with gaps in the alignment are shown in gray. **B)** Predicted structure of the *A. mellifera* Mlh1 protein from AlphaFold 2. Regions are colored by the confidence of the prediction (pLDDT score). Amino acids affected by missense SNP:s are marked in green and labeled with their positions.
